# An updated and spatially validated somatic single-cell atlas of *Hydractinia symbiolongicarpus*

**DOI:** 10.1101/2025.06.03.657738

**Authors:** Jingwei Song, Danielle de Jong, Justin Waletich, Andreas D. Baxevanis, Christine E. Schnitzler

## Abstract

Single-cell RNA sequencing (scRNA-seq) has revolutionized transcriptomic research, enabling the creation of detailed tissue, organ, and species-level atlases for model organisms. In *Hydractinia*, a cnidarian model for stem cell and regeneration studies, recent atlases have revealed key insights into cell types and developmental processes. However, these atlases remain limited in cell numbers and transcriptomic depth and cell type assignments were largely made *in silico*. Here, we present an updated *Hydractinia* single-cell atlas by integrating new datasets from fixed cells with previously published live-cell data. This expanded atlas captures over 47,000 cells from feeding polyps and stolon tissue, recovering and refining major somatic cell lineages including cnidocytes, neurons, gland cells, epithelial cells, and stem cells (i-cells), as well as identifying a novel population of putative immune cells. We investigated the spatial expression patterns of selected marker genes and validated all major cell types and several cell states. Our analyses uncovered a previously undescribed neural subtype, two spatially distinct gland cell populations, a stolon-specific cell type, and a putative immune cell cluster. Additionally, we recovered and explored a complete *Hydractinia* cnidocyte trajectory with two distinct endpoints, supported by spatial marker gene expression that reflects the developmental progression of cnidoblasts as they mature and migrate towards the tentacles. Subclustering of somatic i-cells revealed putative progenitor states and a potential population of true stem cells. Together, this atlas significantly advances our understanding of *Hydractinia* cellular diversity and dynamics, allowing us to generate new hypotheses and provide a valuable resource for the cnidarian research community and beyond.

## Introduction

*Hydractinia* is a fascinating genus of colonial marine hydrozoans that has captivated scientists since the late 19th century, largely because of its remarkable stem cell biology and extraordinary regenerative abilities, allowing the organism to regrow any part of its body at any time (Weismann 1883). These organisms have experienced a resurgence of interest in the molecular era owing to their suitability for genetic manipulation, microscopy, and molecular studies (Künzel et al. 2010; Frank et al. 2020). Advances in genomics, including the sequencing of new genomes, transcriptomes, and the development of transgenic lines, have positioned *Hydractinia* as an emerging model organism for studying fundamental biological processes including regeneration and stem cell biology.

The *Hydractinia* colony consists of multiple polyp types connected via a basal mat encompassing a network of endodermal gastrovascular canals called stolons, located between two ectodermal epithelial layers. The two most common polyp types in cultured animals are gastrozooids (feeding polyps) and gonozooids (sexual polyps). In nature, defensive polyps are also present (dactylozooids and tentaculozooids). The feeding polyp has ectodermal (epidermal) and endodermal (gastrodermal) layers that are separated by an acellular mesoglea; it also contains several specialized cell types such as neurons, cnidocytes, epithelial cells, and gland cells. *Hydractinia* also contains a population of stem cells called interstitial cells (i-cells; Fig. 1a). These i-cells have traditionally been characterized by their size, morphology, location, and staining properties (Müller 1964; Müller 1967; Plickert et al. 1988) and, more recently, through the expression of marker genes such as *Piwi1* and *Vasa* (Rebscher et al. 2008; Plickert et al. 2012). However, it remains unclear whether the cell populations identified using different methods (e.g., morphology, staining, or marker gene expression) represent the same cellular population or if these methods also include progenitor populations. *In situ* hybridization (ISH) studies of various stem cell marker genes have generally confirmed the locations of i-cells in the colony but have revealed discrepancies in the number and morphology of cells stained, suggesting potential heterogeneity within these populations (Bradshaw et al. 2015; Waletich et al. 2024). Single-cell RNA sequencing holds great promise for resolving these questions, as it allows for the bioinformatic clustering of cells with overlapping transcriptomic profiles and the identification of cell-type marker genes, providing insights into cellular diversity, cell states, and function.

**Figure 1.**
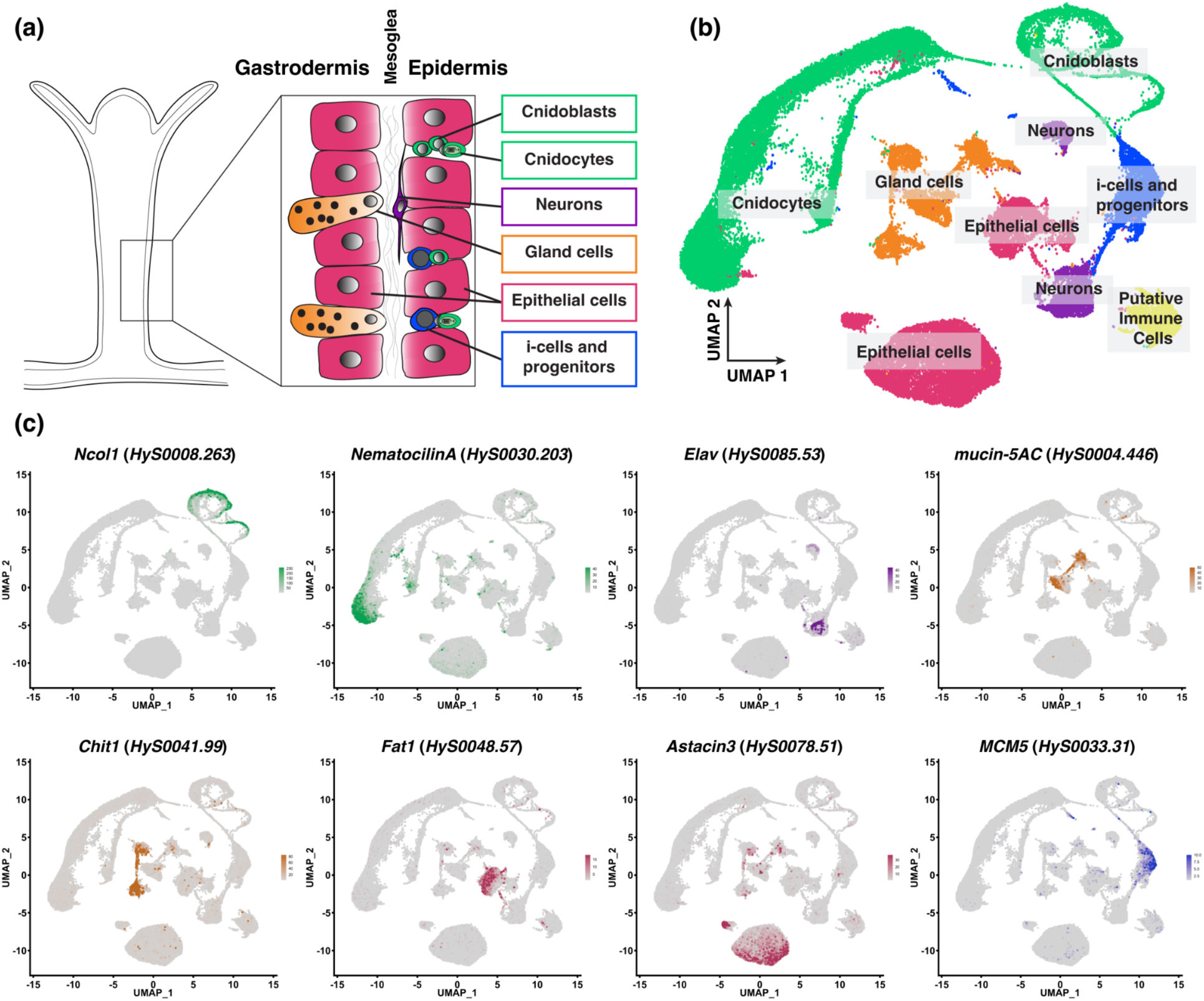
Overview of *Hydractinia* feeding polyp cell types and the updated single-cell atlas. (a) Schematic of a feeding polyp, showing the major cell types present in the two cell layers (gastrodermis and epidermis), separated by the mesoglea. (b) Two-dimensional Uniform Manifold Approximation and Projection (UMAP) representation of the updated *Hydractinia* single-cell atlas (47,901 cells), with major cell states and cell types labeled. (c) UMAP expression of specific genes that characterize the different cell states/cell types. Colors are consistent between all panels: green indicates cnidoblasts and cnidocytes, purple indicates neurons, orange indicates mucous and zymogen gland cells, maroon indicates ectodermal and endodermal epithelial cells, dark blue indicates i-cells and progenitors, and yellow indicates the putative immune cells.

The advent of single-cell RNA sequencing (scRNA-seq) has revolutionized genomics by enabling researchers to analyze transcriptomes of individual cells (Macosko et al. 2015). Initially applied to mammalian models, this technology has since been utilized to study a wide range of organisms, including various cnidarian species (Sebé-Pedrós et al. 2018; Siebert et al. 2019; Chari et al. 2021; Levy et al. 2021; Steger et al. 2022; Hu et al. 2023). Two previous studies using two different approaches generated single-cell atlases for *Hydractinia symbiolongicarpus* (hereafter, *‘Hydractinia’,* unless specified otherwise), and provided many new biological insights, including one study that combined cells from feeding polyps, sexual polyps, and stolons and provided a somatic i-cell cluster, as well as a germ i-cell cluster that was connected to a complete trajectory of spermatogenesis (Schnitzler et al. 2024), and another study that investigated the distribution of cell types across the colony by separately profiling stolons and two polyp types (Salamanca-Díaz et al 2025). These efforts also faced limitations such as being based on a relatively low number of cells, leading to several small cell clusters that were difficult to characterize (Schnitzler et al. 2024) or limited gene coverage per cell, leading to shallow transcriptomic depth that resulted in unresolved and overlapping clusters that were not spatially validated (Salamanca-Díaz et al. 2025). These atlases can be significantly improved by increasing both the number of cells analyzed and the depth of gene coverage, resulting in a more comprehensive and higher-resolution single-cell atlas followed by spatial validation of cell clusters. This, in turn, will provide deeper biological insights, facilitate further hypothesis generation, and serve as a valuable resource for future studies.

Schnitzler et al. (2024) identified two distinct subpopulations of i-cells in their *Hydractinia* single-cell atlas: one that gives rise to germ cells and another that differentiates into either cnidoblasts (stinging cell progenitors) or neurons. However, the low number of i-cells captured in that study limited further exploration of these i-cell subpopulations. This new study aims to overcome the limitations from previous studies by providing an improved, comprehensive scRNA atlas that encompasses all somatic cell lineages in *H. symbiolongicarpus*. This new atlas has allowed for the identification of cell subtype clusters, including multiple cnidocyte, neural, gland, and epithelial cell clusters, with subsequent spatial validation of these clusters using cell-type-specific molecular markers. We have also performed subclustering analysis of the i-cell population to further understand the structure and transcriptional dynamics of the overall i-cell and progenitor cell populations.

Here, we present an updated *Hydractinia* single-cell atlas containing over 47,000 cells derived primarily from feeding polyp and stolon tissue from the male 291-10 strain, with the atlas being comprised of 19 cell-type and cell-state clusters. We have validated all major cell types in these clusters, as well as select cell differentiation states, using fluorescent *in situ* hybridization (FISH) and hybridization chain reaction-FISH (HCR-FISH) methodologies. This approach has revealed several major findings, including a previously undescribed neural subtype, two spatially separate gland cell populations resembling those found in *Hydra*, a stolon-specific cell type and a putative immune cell cluster. In addition, we provide the first complete cnidocyte trajectory for *Hydractinia* and have validated expression of several markers along this trajectory. Finally, we subclustered the somatic i-cell cluster and were able to identify and assign putative cell states to i-cell subclusters. This subclustering analysis revealed a potential population of true i-cells as well as early progenitor populations. We have identified multiple cell-type and cell-state specific markers that can be used to investigate biological phenomena such as regeneration and cellular differentiation. These markers also reveal previously unrecognized transcriptional diversity within specific cell types that may underlie functional differences within the animal, allowing us to gain a deeper understanding of the genetic mechanisms governing the differentiation of progenitor cells into specific cell types. Additionally, we identify candidate markers whose regulatory sequences can be used to drive fluorescent reporters in a cell type- or cell state-specific manner.

## Results and Discussion

We initially processed our raw sequencing reads using Cell Ranger v7 (Supplementary Table S1) and found that two previously generated live-cell libraries from feeding polyp, sexual polyp, and stolon tissue exhibited the highest median genes per cell (792), fraction of reads in cells (83.3%), and sequencing saturation (92.6%; Schnitzler et al. 2024). Our new data were derived exclusively from feeding polyp and stolon tissue, using fixed samples (methanol and ACME). The methanol-fixed dataset, obtained from two libraries, had the highest mean number of total genes detected (17,786) but the lowest fraction of reads in cells (33%). The ACME-fixed dataset from seven libraries had the highest mean reads per cell (162,991) and a moderate fraction of reads in cells (66%). Although there were some variations in the summary statistics, we considered all libraries to be of adequate quality and proceeded with dataset integration.

In this study, we focused on the somatic cell types of *Hydractinia* and excluded germ cells. For the previously generated live-cell dataset, we first removed mature sperm cells and doublets (n=4,865), which together comprised about 50% of that dataset. After quality control (QC) of each fixed cell dataset and subsequent integration of all three datasets, we obtained a final merged dataset comprised of 47,901 cells (Methanol fixed: 26,247; ACME fixed: 17,631; Live cells: 4,023) (Fig. 1b). By combining known cell-type specific genes with the results of a literature search for the top differentially expressed genes, we identified five major somatic cell lineages in *Hydractinia*: cnidocytes (green), neurons (purple), gland cells (orange), epithelial cells (magenta), and i-cells/progenitors (blue) (Fig. 1b). We also identified a cluster (yellow) that could not be readily assigned to any known cell type but which we have putatively labeled as immune cells, a designation that is discussed below (Fig. 1b, Fig. 1c). A higher-resolution UMAP illustrating the clusters that comprise each cell lineage can be found in Supplementary Fig. S1. A list of diagnostic genes used to annotate the clusters is provided in Supplementary Table S2, and their expression patterns within the single-cell atlas is shown as Supplementary Fig. S2.

### Cnidocytes and Cnidogenesis

One of the most prominent features of the atlas is the developmental trajectory of cnidocytes (cnidogenesis; Fig. 2a). Cnidoblasts (C6, C15, C16) and developing cnidocytes (C4) were intermediary between the i-cell and progenitor cluster (C3) and the mature cnidocytes (C1, C2, C8) (Supplementary Fig. S1). Both CytoTRACE and pseudotime analyses confirmed the maturation of cnidocytes along this trajectory (Fig. 2a). Notably, the single-cell atlas, CytoTRACE, and pseudotime analyses all revealed a single cnidogenesis pathway that splits into two terminal endpoints (Fig. 2a, “1” and “2”). A heatmap generated using a combination of known cnidogenesis genes and novel markers highlighted a subset of genes expressed along this trajectory (Fig. 2b). Spatial expression analysis shows that early-cnidogenesis genes such as *Txd12* (Fig. 2c-c’’’) and *Fkbp14* (Supplementary Fig. S3 a-a’’’), as well as mid-cnidogenesis genes such as *Ncol1* (Fig. 2d-d’’)’ and *Dkk3* (Supplementary Fig. S3 b-b’’’), are all expressed in the lower half of the feeding polyp body column, a region known to harbor i-cells and cnidoblasts (Klompen et al. 2022; Waletich et al. 2024).

**Figure 2.**
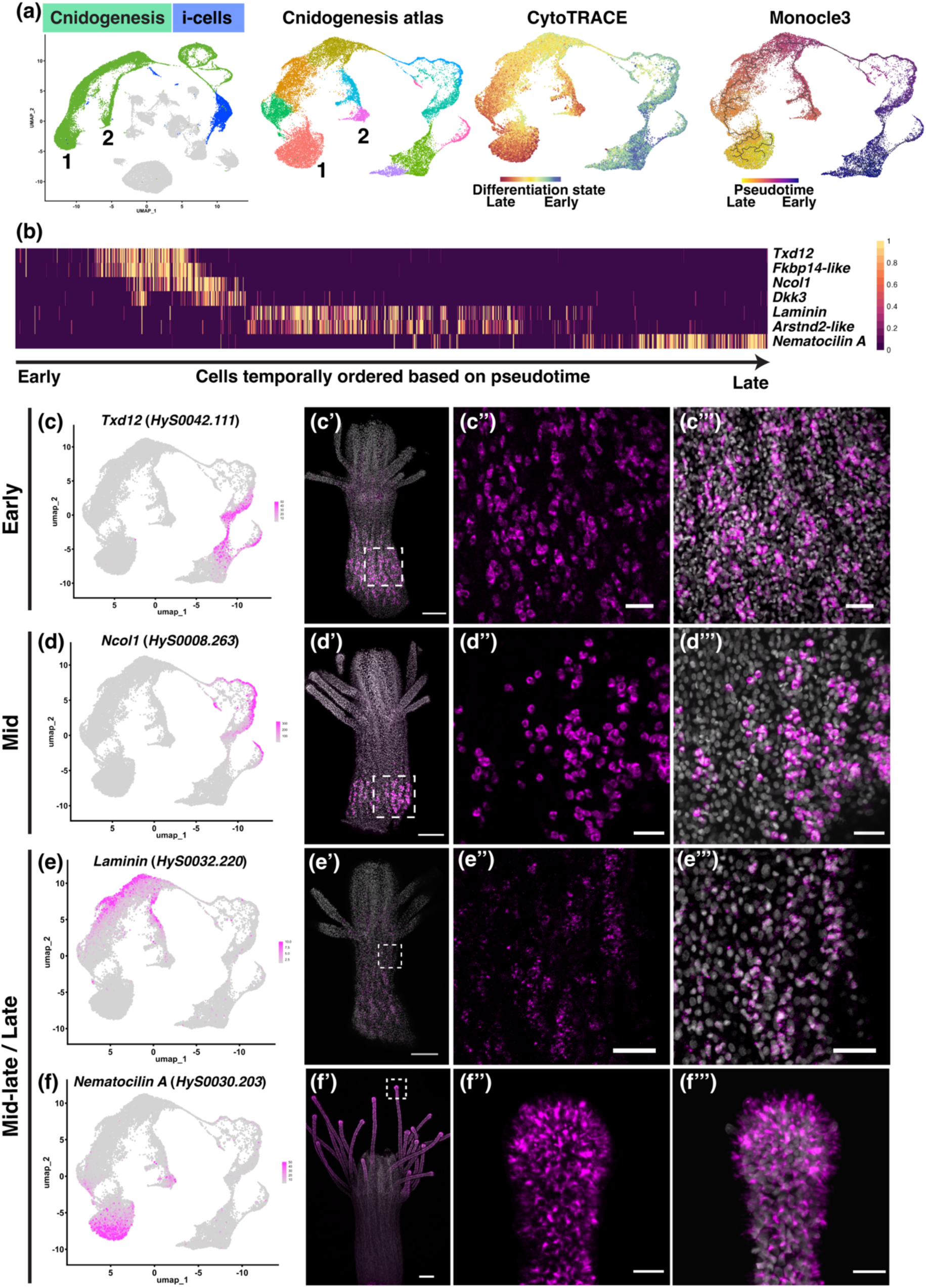
Cnidogenesis trajectory from i-cell to mature cnidocytes. (a) Single-cell atlas showing cells involved in cnidogenesis (green) originating from a single cluster of i-cells/progenitor cells (blue). These clusters were selected and subjected to reclustering to form the cnidogenesis atlas, upon which both CytoTRACE and Monocle3 analyses were performed. CytoTRACE shows the differentiation state from early (blue/green) to late (orange/red), while Monocle3 provides a pseudotime analysis from early (purple) to late (yellow). The two endpoints in the single cell atlas and cnidogenesis atlas are labeled as 1 and 2, representing transcriptionally distinct, fully differentiated cnidocytes. (b) Heatmap depicting the normalized expression of seven selected genes. Each column represents an individual cell that was ordered based on their pseudotime values from the lowest (earliest) to the highest (latest). (c-f’’’) Left-most column shows the expression of a particular gene in the cnidogenesis atlas, while panels to the right show expression of that gene in an adult feeding polyp. Dotted white boxes in (c’), (d’), (e’) and (f’) indicate the regions shown at higher magnification in (c”)-(c’’’), (d”)-(d’’’), (e”)-(e’’’) and (f”)-(f’’’) respectively. Gene expression is shown in magenta and nuclei are shown in grey. *HyS0042.111* (*Txd12*) is a marker of early cnidogenesis (c)-(c”), *HyS0008.263* (*Ncol1*) is a marker of mid-cnidogenesis (d)-(d’’’), while *HyS0032.220* (*Laminin*) and *HyS0030.203* (*Nematocilin A*) are markers of mid-late/late cnidogenesis (e)-(f’’’). Scale bars: 100 µm in (c’), (d’), (e’) and (f’); 20 µm in all other panels.

Mid-to-late stage cnidogenesis genes such as *Laminin* (Fig. 2 e-e’’’) and *Arstnd2-like* (Supplementary Fig. S3 c-c’’’) showed expression in cells in not only the lower half of the body column, but also in cells extending towards the upper half. These genes were often detected in cells with distinguishable cnidocyte capsules, supporting the idea that cnidocytes migrate as they mature (Thomas and Edwards 1991; Tardent 1995; Schnitzler et al. 2024). A late-stage/mature cnidogenesis marker, *Nematocilin A* (*HyS0030.203*) (Hwang et al. 2008) was present at both endpoints of the trajectory (C1/C8, Fig. 1c) and was expressed exclusively in the tentacles (Fig. 2f-f’’’) (Schnitzler et al. 2024), confirming the two end branches of this trajectory (C1 and C8) contain mature cnidocytes. There are two major cnidocyte types known in adult *Hydractinia* feeding polyps: desmonemes and euryteles. Desmonemes are smaller than euryteles, and the morphology of their cnidocyst capsules also differs between the two types (Mills 1976; Lange et al. 1989; Schuchert 2014). Given that desmonemes constitute 75% of tentacle cnidocytes in *Hydractinia* and euryteles constitute the remaining 25% (Klompen et al. 2022) and considering that C1 is larger than C8 and expressed *Hydra* desmoneme markers (Supplementary Table S2), we hypothesized that C1 corresponded to desmonemes and C8 to euryteles. To test this hypothesis, we designed HCR-FISH probes for two new marker genes expressed specifically in each of the two clusters: *HyS0002.425* (at the tip of C1) and *HyS0027.82* (at the tip of C8). *HyS0002.425* was expressed in cells throughout the tentacles and only sparsely in the body column (Supplementary Fig. S3d-d’’’). *HyS0027.82* was expressed in cells concentrated at the tentacle tips, as well as in cells sparsely distributed throughout the tentacles (Supplementary Fig. S3e-e’’). A double HCR-FISH experiment showed that the two populations of cnidocytes were non-overlapping (Supplementary Fig. S3d’’’). *HyS0027.82^+^*cnidocytes appeared larger than *HyS0002.425^+^* cnidocytes in transmitted light images (Supplementary Figs. S3e’’’, f-f’) but the dense packing of cnidocytes in the tentacles made size comparisons somewhat difficult. Overall, the results supported our hypothesis that the larger end branch represents desmonemes (C1) and the smaller branch represents euryteles (C8).

Comparison of our cnidogenesis trajectory with those of other cnidarians revealed a close resemblance to that of *Hydra* (Cazet et al. 2023), where mature cnidocytes form distinct clusters based on their type. This contrasts with findings from *Nematostella vectensis*, where the cnidocyte trajectory derives from a single pool of progenitors that splits into multiple differentiation pathways corresponding to the different cnidocyte types before the trajectory converges into a single cluster (Steger et al. 2022; Cole et al. 2024). Such contrasting trajectories raise questions about the evolution of cnidocytes and cnidocyte types in different cnidarian taxa, which is beyond the scope of the current study.

### Neurons

Two neural cell types have been described in *Hydractinia echinata*, a sister species to *H. symbiolongicarpus*: sensory neurons and ganglionic neurons. This classification is based primarily on morphological characteristics observed using electron microscopy (Stokes 1974). The two neural cell types can also be distinguished by their orientation relative to the mesoglea, as ganglionic cells lie parallel to the mesoglea, whereas sensory cells are oriented perpendicular to it (Thomas and Edwards 1991). Together, these two neural types form the ectodermal nerve net of *H. echinata* (Stokes 1974). In our cell atlas, we identified two clusters (C7 and C14) with neural gene expression signatures (Fig. 1b, Fig. 3a), comprised of classic neural markers such as *Elav* (*HyS0085.53*, Fig. 1c) (Pascale et al. 2008; Nakanishi et al. 2012) and *Neurocalcin* (*HyS0034.90*) (Vijay-Kumar and Kumar 2002) (Supplementary Fig. S2). The neuropeptide precursor genes for *RFamide* (*HyS0013.338*) and *GLWamide* (*HyS0009.155*) were expressed exclusively in C7. The *achaete-scute homolog* (*Ash, HyS0005.437*) was expressed in C14 and in developing cnidocytes (C4), similar to its expression in *Hydra magnipapillata*, where it is a known marker of sensory neurons and differentiating cnidocytes (Hayakawa et al. 2004). To determine the spatial location of cells constituting the two clusters of *Hydractinia* neurons (“Neurons A” and “Neurons B”, Supplementary Fig. S1), we performed double HCR-FISH using the gene encoding for *RFamide* (*HyS0013.338*) as a marker of C7 cells (Fig. 3b) and *HyS0049.55*, a putative neuropeptide (see below) as a marker of C14 cells (Fig. 3c). Consistent with previously published results (Schmich et al. 1998; Chrysostomou et al. 2022), *RFamide^+^* cells were predominantly located in the hypostome, and were present in the body column and tentacles, albeit less densely. In contrast, *HyS0049.55^+^* cells were predominantly located in the tentacles, revealing a previously undescribed population of neurons in the *Hydractinia* feeding polyp (Fig. 3d-f’’’). We did not observe any overlap in the expression of *RFamide* and *HyS0049.55*. Based on our double HCR-FISH experiments, we were unable to determine whether Neurons A and Neurons B correspond to ganglionic and sensory types based on cell orientation or other spatial information. It is possible that one or both of these two neuron types represent a mixture of ganglionic and sensory neurons, as previous studies in *Hydractinia* have indicated that both types are present in the hypostome (Klimovich et al. 2018; Chrysostomou et al. 2022). To further investigate the morphology of the neurons expressing each of our marker genes, we performed HCR-FISH on dissociated cells (Fig. 3g-l’). We detected a range of morphologies, including tripolar neurons (Fig. 3g-g’, j-j’), unipolar neurons (Fig. 3h-h’, k-k’), and bipolar neurons (Fig. 3i-i’, l-l’) However, there was no clear distinction in the morphology of neurons from C7 and C14. Further study will be required to resolve which morphological types are present in each neuron cluster.

**Figure 3.**
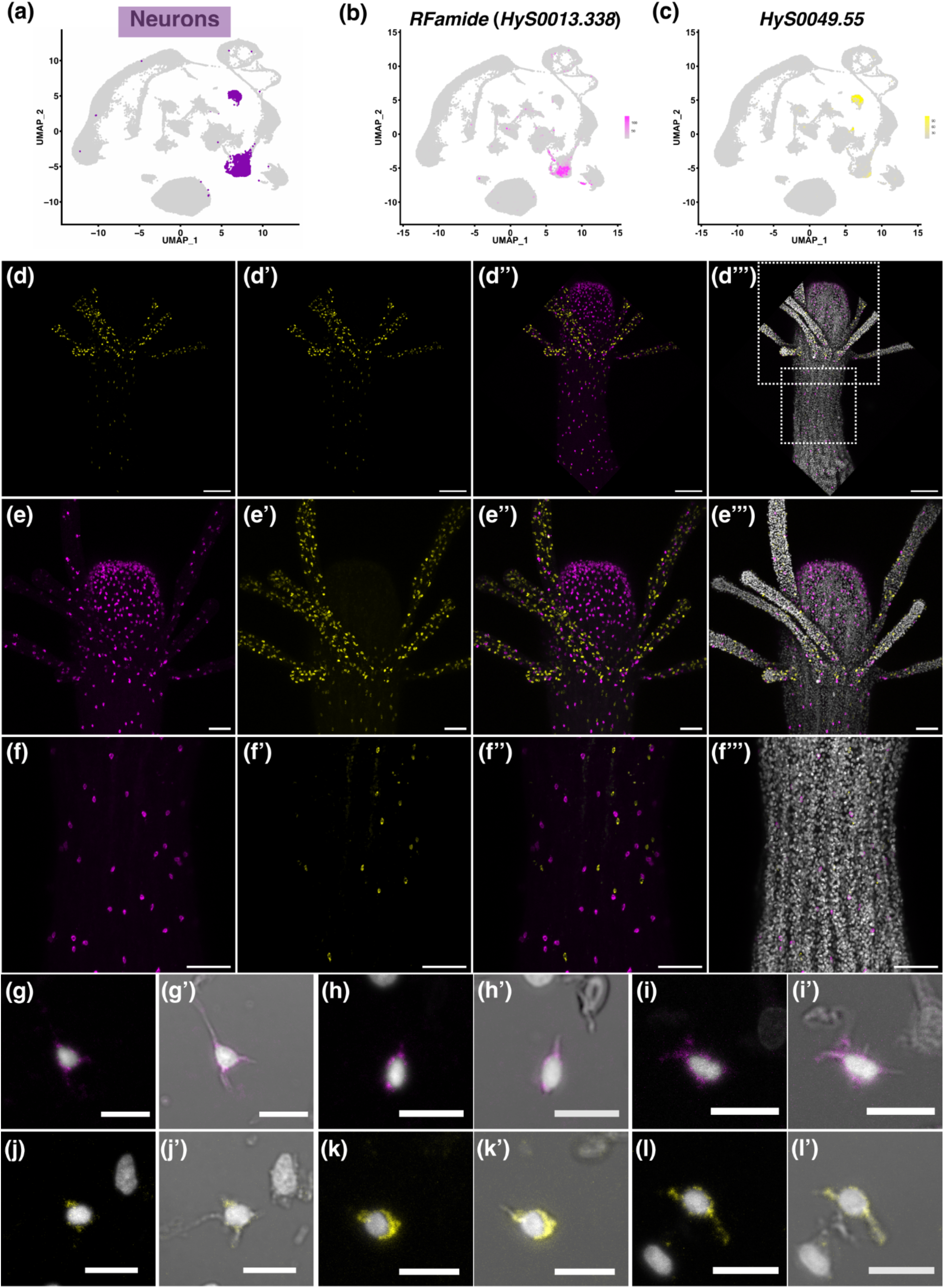
Expression analysis of markers expressed in differentiated neuron cell clusters. (a) Two-dimensional UMAP of the *Hydractinia* single-cell atlas, with neuron clusters highlighted in purple. (b) UMAP showing expression of *RFamide* (*HyS0013.338*), which specifically marks a subset of cells in cluster 7 (magenta; this color also marks *HyS0013.338*^+^ cells throughout the figure). (c) UMAP showing expression of *HyS0049.55*, which specifically marks the majority of cells in cluster 14 (yellow; this color also marks *HyS0049.55*^+^ cells throughout the figure). (d)-(f’’’) Confocal images of HCR-FISH of the genes shown in (b) and (c) in adult feeding polyps. The white dotted boxes shown in (d’’’) indicate regions selected for higher magnification images in (e)- (e’’’) and (f)-(f’’’). Nuclei are shown in grey. (g)-(l’) Cell dissociations followed by HCR-FISH show a range of neural morphologies. Scale bars: 100 µm in (d)-(d’’’), 50 µm in (e)-(f’’’), and 10 µm in all other panels.

The nervous system of cnidarians is thought to be primarily governed by peptidergic signaling, with neuropeptides playing a role in many aspects of its biology, including metamorphosis, specific behaviors, reproduction, and feeding (Leitz et al. 1994; Grimmelikhuijzen et al. 1996; Takeda et al. 2018; Attenborough et al. 2019; Takahashi 2020; Weissbourd et al. 2021; Yamamoto and Yuste 2023). As there were no obvious morphological differences between the cells expressing our C7 and C14 markers, we sought to identify the complement of neuropeptides expressed in each cluster, with the goal of identifying potential functional differences between the two neural clusters, A similar approach was taken by Chari et al. (2021) in a study of the hydrozoan jellyfish *Clytia*. Using sequence-based analyses, we identified 12 putative neuropeptides, including two that were previously known (*RFamide* and *GLWamide*) and 10 that were previously unidentified (Supplementary Fig. S4, Supplementary Table S3). The coding sequence of one of these newly identified neuropeptides (*HyS0049.55*) was used as a C14-specific marker in experiments described above. Some of these novel neuropeptides appear to be related to those previously isolated from cnidarians. For example, *HyS0052.141* shows similarity to the PRXamide family of neuropeptides that are present in many invertebrates, including cnidarians, and specifically to the maturation inducing hormones (MIH; RPRAmide peptides). MIH have been shown to be synthesized directly by cells in the gonad in two hydrozoan jellyfish and to act directly in oocyte maturation (Takeda et al. 2018). Other putative neuropeptides identified do not show clear similarity to previously described neuropeptides. The neuropeptide precursors we identified were present in one or both of the neural clusters in the single-cell atlas. This phenomenon, in which distinct combinations of neuropeptides are produced by different populations of neurons, has been observed previously, specifically in studies of the nervous system in both *Hydra* and *Clytia,* and is thought to be related to functional differences (Grimmelikhuijzen et al. 2002; Chari et al. 2021; Yamamoto and Yuste 2023; Prabhu and Reddy 2025). Future in-depth analyses of neural location, neurochemistry (including the presence of classical chemical neurotransmitters), and spatial and functional analyses of neurons in different polyp and tissue types within the *Hydractinia* colony will be required to fully elucidate the diversity of neuron types and neural functions in this animal.

### Gland cells

The characterization and distribution of the different types of gland cells in *Hydractinia* polyps have not been well-studied. Studies in *Hydra* have identified two broad types of gland cells: mucous gland cells (having two subtypes, spumous and granular) that secrete mucus, and zymogen gland cells that secrete proteolytic enzymes into the gastric cavity to enable digestion of food particles (Rose and Burnett 1968a; Haynes and Davis 1969). Based on selected markers of genes known to be expressed in gland cells in *Hydra* and *Hydractinia* (Augustin et al. 2006; Schwarz et al. 2007; Siebert et al. 2019; Cazet et al. 2023; Schnitzler et al. 2024), we annotated four clusters in the single-cell atlas as corresponding to the two types of gland cells: mucous gland cells (clusters C10 and C11) and zymogen gland cells (clusters C12 and C13) (Fig. 4a, Supplementary Fig. S1, Supplementary Table S2).

**Figure 4.**
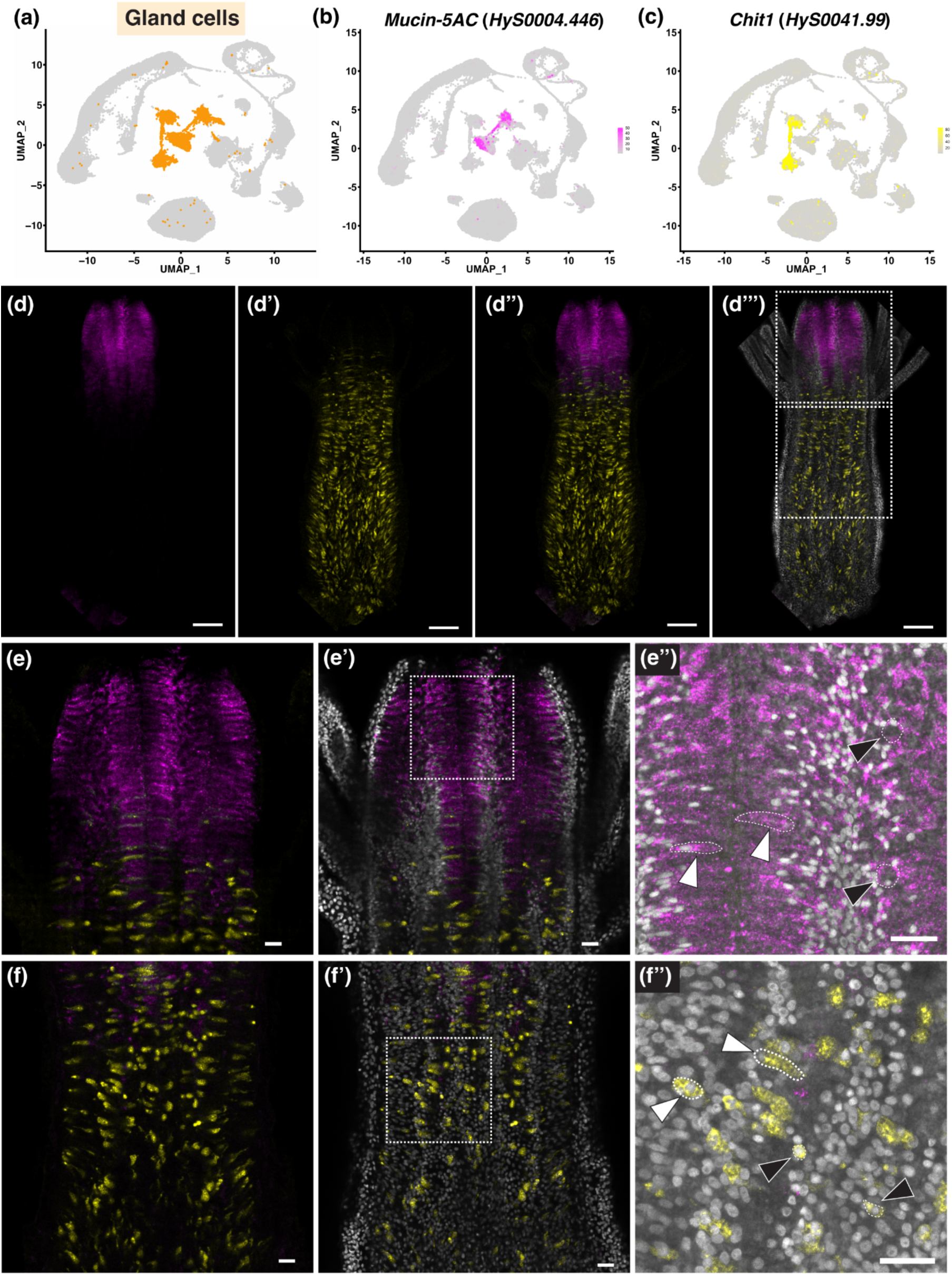
Expression analysis of markers expressed in differentiated gland cell clusters. (a) Two-dimensional UMAP of the *Hydractinia* single-cell atlas with gland cell clusters highlighted in orange. (b) UMAP showing expression of the mucous gland cell marker *mucin-5AC* (*HyS0004.446*) (magenta; this color also marks *HyS0004.446*^+^ cells throughout this figure). (c) UMAP showing expression of the zymogen gland cell marker *Chit1* (*HyS0041.99*) (yellow; this color also marks HyS0041.99^+^ cells throughout this figure). (d)-(f”). Confocal slices of an HCR-FISH in the gastroderm of adult feeding polyps, showing expression of the genes shown in (b)-(c). Nuclei are shown in grey. (d)- (d’’’) shows a whole adult feeding polyp, while (e)-(e’’) shows a higher magnification image of the hypostome and (f)-(f’’) shows a higher-magnification image of the polyp body. Dotted white boxes in (d’’’) indicate the regions of higher magnification images shown in (e-e’’) (upper white dotted box) and (f-f’’) (lower white dotted box). Dotted white boxes in (e’) and (f’) indicate the regions shown at higher magnification in (e”) and (f”), respectively. Images in (e’’) and (f’’) are confocal slices overlaid with transmitted light images. White arrowheads in (e”) indicate examples of long, elongated *mucin-5AC*^+^ cells, while black arrowheads in (e”) indicate smaller, rounder *mucin-5AC*^+^ cells. White arrowheads in (f”) indicate examples of large *Chit1*^+^ cells with intracellular granules, while black arrowheads indicate examples of small, rounded *Chit1*^+^ cells. Scale bars: 100 µm in (d)-(d’’’) and 20 µm in all other panels.

To determine the relative spatial locations and cellular morphology of the cells in the putative mucous and zymogen gland cell clusters and confirm their annotation, we selected a marker exclusive to the putative mucous gland cells clusters (C10 and C11), *mucin-5AC* (*HyS0004.446,* Fig. 4b), and performed double HCR-FISH together with a previously validated gene marker for zymogen gland cells (C12 and C13), *Chitinase 1* (*Chit1, HyS0041.99)* (Schnitzler et al. 2024) (Fig. 4c).

Expression analyses showed that many tightly packed *mucin-5AC*^+^ cells were present in the gastroderm of the hypostome, while *Chit1*^+^ cells were distributed as expected throughout the gastroderm of feeding polyp bodies, aboral to the hypostome (Fig. 4d-f’’; (Schnitzler et al. 2024). The two populations – *mucin-5AC*^+^ and *Chit1*^+^ cells – were mostly spatially separate, except for a region at the base of the tentacles where they were adjacent, with some intermixing of cell types (Fig. 4e). Spatial expression analysis confirmed our assignment of C10 and C11 cells as mucous gland cells and C12 and C13 as zymogen gland cells. While there are differences among hydrozoan species regarding the distribution of mucous and zymogen gland cells in the polyp body, the most well-studied *Hydra* species contains only mucous gland cells in the gastrodermis of the hypostome and only zymogen gland cells in the gastrodermis of the polyp body (Rose and Burnett 1968a; Haynes and Davis 1969; Siebert et al. 2008), consistent with our observations in *Hydractinia* feeding polyps. This spatial separation of gland cell types was also previously reported for *Hydractinia echinata* via transmission electron microscopy (Thomas and Edwards 1991).

The *mucin-5AC*^+^ cells appear to exhibit two different morphologies, but their tight packing in the hypostome makes clear descriptions challenging. In general, we observed round cells with what appears to be large intracellular vacuoles (Fig. 4e’’ black arrowheads) and long, thin cells (Fig. 4e’’ white arrowheads). In *Hydra,* two types of mucous gland cells are present (spumous mucous and granular mucous gland cells) that are morphologically and functionally distinct (Siebert et al. 2008; Siebert et al. 2019; Cazet et al. 2023), so it is possible that the two morphological variants seen in *Hydractinia* correspond to these subtypes. *Chit1*^+^ zymogen gland cells were not as tightly packed together in the gastrovascular cavity and had distinct cell boundaries, allowing us to more confidently identify the two distinct morphologies we observed. Cells were either small and round/oval (approximately 10 µm in diameter) with large nuclei (Figure 4f’’, black arrowheads), or large (approximately 15-20 µm in length) with intracellular vacuoles or granules (Fig. 4f’’, white arrowheads). The appearance of the larger *Chit1*^+^ cells is consistent with the known morphology of mature zymogen gland cells. The smaller *Chit1*^+^ cells resemble a cell type observed in the gastrodermis of the hydrozoans *Halocordyle disticha* and *Hydra viridissima*, where they have been termed “young zymogen cells”, “undifferentiated gastrodermal cells”, or “basal reserve cells” (Bouillon, J. 1966; Rose and Burnett 1968b; Haynes and Davis 1969; Thomas and Edwards 1991). These cells were described as oval in shape, 10-12 µm in diameter, and were hypothesized to be either immature zymogen gland cells, a zymogen gland cell following secretion of their granules, or dedifferentiated zymogen gland cell (Rose and Burnett 1968b; Haynes and Davis 1969). In the gastroderm of the stolons and budding feeding polyps in young *Hydractinia* colonies, we have observed that almost all *Chit1*^+^ cells are of this smaller type (unpublished data), suggesting that these cells might indeed be precursors of mature zymogen gland cells.

Our single cell atlas has identified four clusters of gland cells and shows that they are comprised of mucous and zymogen gland cells, the two major gland cell types found in *Hydractinia*. Spatial expression analyses using markers of both of these cell types show that, as in *Hydra*, these cells are located exclusively in the gastrodermis and are spatially separated along the oral-aboral axis: mucous gland cells are exclusively found in the hypostome, while zymogen gland cells are found exclusively in the body column, with only a small amount of mixing at the boundary of these two regions. Further investigations are required to determine whether the two clusters of zymogen gland cells and the two clusters of mucous gland cells correspond to different subtypes. It is possible that the two mucous gland cell clusters correspond to spumous and granular mucous gland cells as described in *Hydra* (Siebert et al. 2008). For example, *mucin2* (*HyS0015.116*) is restricted to cluster 10, and its probable ortholog in *Hydra* is specifically expressed in spumous mucous gland cells (G010426; Cazet et al. 2023). Alternatively, the different clusters of gland cells could correspond to the different cellular morphologies we observed in our spatial gene expression analyses, to differences in function, or perhaps to a combination of these factors. *Rhamnospondin* (*HyS0004.396*) is specifically expressed in one of the mucous gland cell clusters (C11) (Supplementary Fig. S2) and most likely plays a role in immune recognition (Schwarz et al. 2007), suggesting a functional difference between cells in C10 and C11. Conducting multi-color spatial expression analyses with markers exclusive to each of the four gland cell clusters will allow us to further investigate these clusters and their contributions to both zymogen and mucous gland cell populations in *Hydractinia* feeding polyps.

### Epithelial cells

Cnidarian epithelial cells are known to be multifunctional (Buzgariu et al. 2015; Leclere and Röttinger 2017; Holstein 2023) and many names have been used in the past to refer to subtypes, such as epitheliomuscular cells (EMCs) (Stokes 1974; Weis and Buss 1987; Leclere and Röttinger 2017) and digestive muscular cells (Thomas and Edwards 1991). We defined the two major epithelial cell clusters based solely on tissue layer location: endodermal (C0 and C17) and ectodermal (C5), while acknowledging their multifunctionality (Fig. 1b, Fig. 5a). A marker for C0 and C17, *Astacin 3* (*HyS0078.51*, Fig. 5b) (Möhrlen et al. 2006), was expressed in the endoderm along the entire polyp body, including in the endodermal cells of the tentacles (Fig. 5d-f’’’). A specific marker for C5, *Fat 1* (*HyS0048.57*, Fig. 5 c), was expressed in cells in the ectoderm along the entire body of the feeding polyp, as well as in the tentacles (Fig. 5d-f’’’). Ectodermal epithelial cells have been shown to have more prominent myofibrils than the endodermal epithelial cells in *Hydractinia* and were hypothesized to be the main drivers of muscular contraction (Dandar-Roh et al. 2004). In support of this hypothesis, we found more cells that expressed muscle-related genes in C5 compared to C0 and C17, such as the genes encoding for the myosin heavy chain structural protein (*HyS0006.325*) and the myosin light chain kinase (*HyS0028.60*). In contrast, the most significant GO term for C0 was “GO:0006508, proteolysis” (Supplementary Fig. S5a), consistent with previous descriptions of a digestive function for gastrodermal epithelial cells (Thomas and Edwards 1991). Notably, many genes in the major endodermal epithelial cluster (C0) showed a gradient expression pattern within the cluster (e.g. *HyS0008.375, HyS0021.141, HyS0034.215*, *HyS0087.37* and *HyS0244.2*) (Supplementary Fig. S5b). Similarly, in *Hydra,* some epithelial cell marker genes displayed graded patterns depending upon their position along the oral-aboral axis (Siebert et al. 2019). Further *in situ* hybridization experiments with these and other genes will be needed to confirm the hypothesis that the graded expression level of genes in this cluster is related to spatial expression patterns along the oral-aboral axis in feeding polyps.

**Figure 5.**
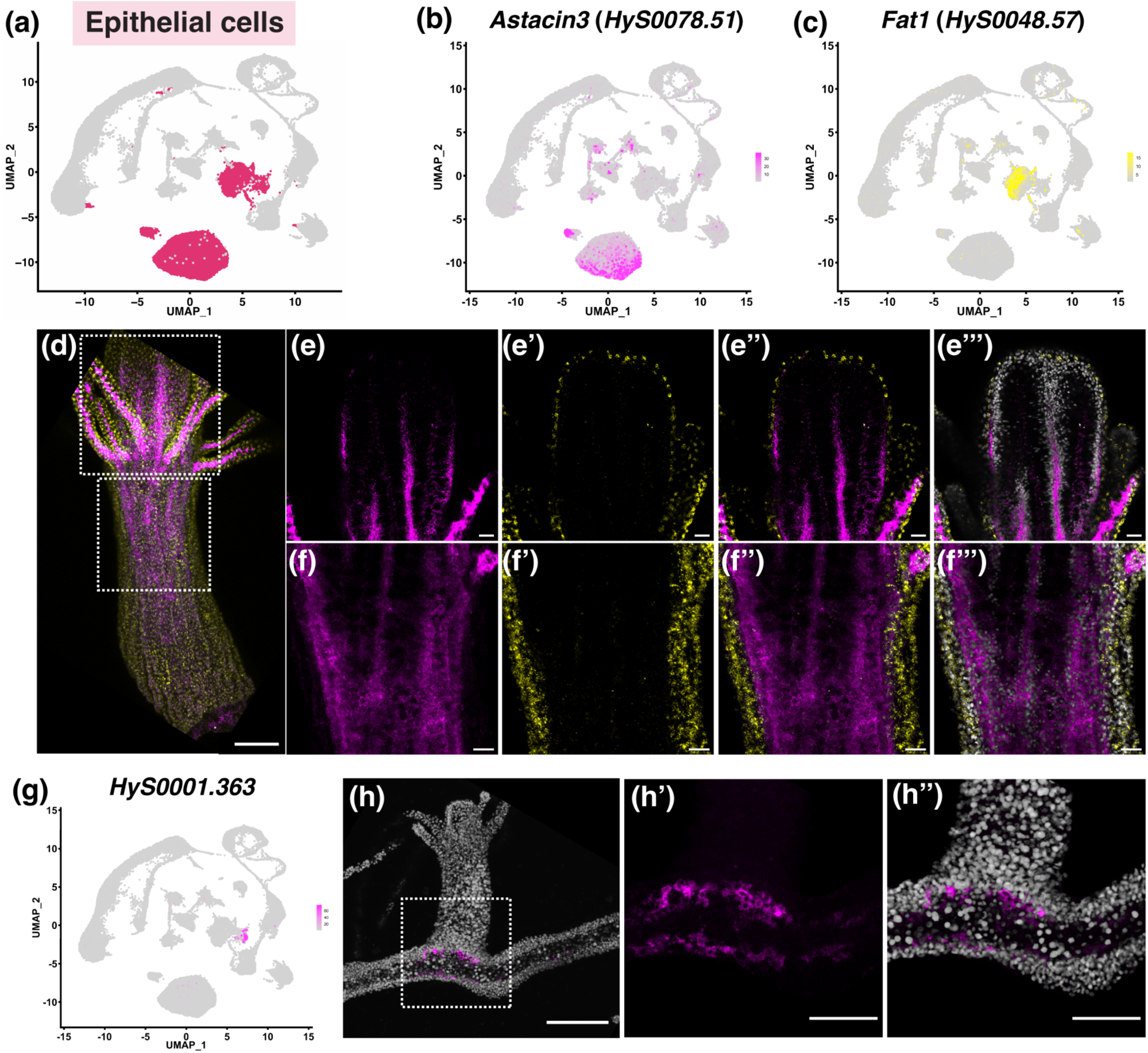
Expression analysis of markers expressed in epithelial cell clusters. (a) Two-dimensional UMAP of the *Hydractinia* single-cell atlas with epithelial cell clusters highlighted in maroon. (b) UMAP expression of *Astacin3* (*HyS0078.51*) highlighted in magenta that marks endodermal epithelial cells. (c) UMAP expression of *Fat1* (*HyS0048.57*) highlighted in yellow that marks ectodermal epithelial cells. (d)-(f’’’) Confocal sections of *in situ* hybridization patterns in an adult feeding polyp of the genes shown in (b)-(c). *Astacin3* (*HyS0078.51*) expression is shown in magenta and *Fat1* (*HyS0048.57*) expression is shown in yellow. Nuclei are shown in grey. (d) Maximum projection of confocal slices of an adult feeding polyp. White dotted boxes indicate the location of the higher magnification images shown in (e)-(e’’’) (upper box) and (f)-(f’’’) (lower box). (g) UMAP expression of *HyS0001.363* that marks stolon-specific ectodermal epithelial cells highlighted in magenta. (h)-(h’’) Expression of HyS0001.363 in the stolon at the base of a young feeding polyp shown in magenta. Nuclei are shown in grey. The white dotted box in (h) represents the region of the higher magnification image selected for (h’)-(h’’). Scale bars: 100 µM in (d), 20 µM in (e)-(f’’’’), 100 µm in (h), and 50 µm in (h’)-(h’’).

We also identified a cluster adjacent to C5 in the atlas (C18) that appeared to be a subtype of ectodermal cells (Supplementary Fig. S1). Among the differentially expressed genes in this small cluster was *frizzled 3* (*HyS0103.16*), a gene previously reported to be expressed only in stolons (Supplementary Table S4, Supplementary Fig. S2) (Sanders et al. 2020). In addition, *chitin synthase* (*HyS0024.60*) was also highly expressed in C18. Chitin is a major component of the periderm that covers *Hydractinia* stolon tissue, but not polyps (Lange and Müller 1991; Frank et al. 2020). This led us to hypothesize that the cells in cluster C18 are stolon-specific epithelial cells. To confirm this, we conducted HCR-FISH with a gene of unknown function that is highly specific to C18, *HyS0001.363* (Fig. 5g), and found it was specifically expressed at the base of young polyps where they intersected with the stolon (Fig. 5h-h’’). Another recent *Hydractinia* single-cell study also identified a stolon-specific epithelial cell cluster (Salamanca-Díaz et al. 2025), and we cross-checked several stolon-specific markers from their cluster with ours, finding a high level of similarity. For example, the homolog of *HyS0001.363 - LOC130636562 -* is specifically expressed in the stolon-specific cluster in the Salamanca-Díaz atlas, and the best match to the highly repetitive *prisilkin/shematrin-like* genes identified in the Salamanca-Díaz atlas is *HyS0026.224*, which is specific to C18 in our atlas. Further investigation of the origin of these stolon-specific cells during *Hydractinia* metamorphosis and development, as well as analysis of the evolutionary conservation of the genes specifically expressed in this cluster, may lead to a greater understanding of the evolution of coloniality.

### Putative Immune Cells

The immune gene repertoire of *Hydractinia* was predicted to be large but no immune cell types have been described to date (Zárate-Potes et al. 2019). In an initial attempt to assign a putative function to cluster 9, we performed GO enrichment analysis. The top GO term was “peptidyl-tyrosine dephosphorylation” (GO:0035335) (Fig 6 b). Over half of the tyrosine phosphatases encoded in the human genome are expressed by immune cells (Mustelin et al. 2005), which led us to suspect immune related functions of this cluster.

**Figure 6.**
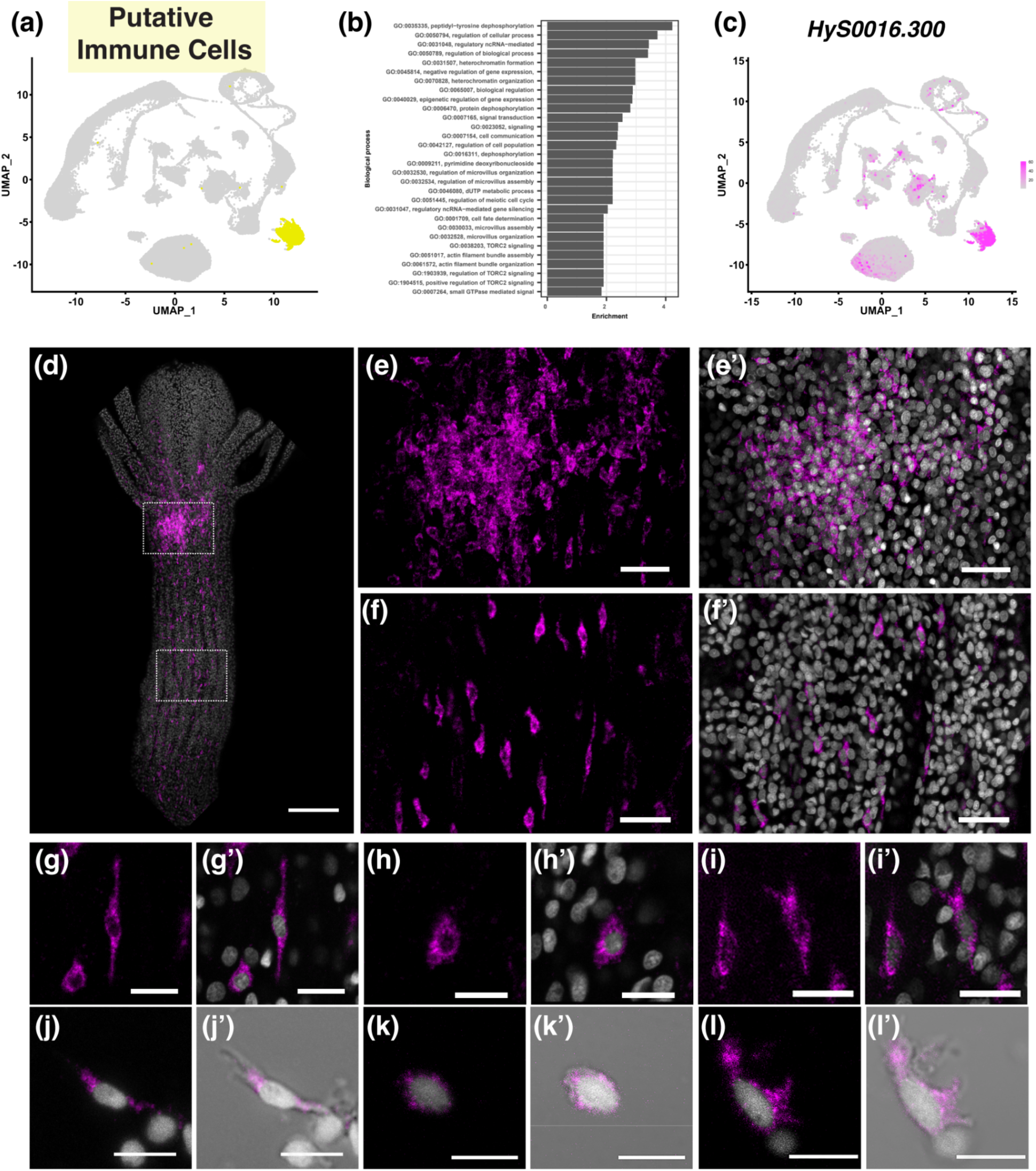
Expression analysis of markers expressed in cluster 9, which are putative immune cells. a) Two-dimensional UMAP of the *Hydractinia* single-cell atlas with cluster 9 highlighted in yellow. (b) Top GO terms associated with differentially expressed marker genes for cluster 9. (c) UMAP expression of *HyS0016.300* highlighted in magenta. (d) Maximum projection of confocal sections of a whole adult feeding polyp. Dotted white boxes show regions of higher magnification images shown in (e)-(e’) (upper box) and (f)-(f’) (lower box). *HyS0016.300*^+^ cells are shown in magenta and nuclei are shown in grey. (g)-(i’) High-magnification images of confocal sections of *HyS0016.300*^+^ cells from an adult feeding polyp illustrating different cell morphologies. (j)-(l’) Cell dissociation followed by HCR-FISH also reveals a range of *HyS0016.300*^+^ cell morphologies. Scale bars: 100 µm in (d), 20 µm in (e)-(f’), and 10 µm in all other panels.

Exploring the list of differentially expressed genes reveals further intriguing hints as to the potential function of the cells in this cluster. For example, one of the top differentially expressed genes is *conodipine-like* (*HyS0053.57*), which contains a phospholipase A2 (PLA2) domain (InterPro entry IPR036444) and a signal peptide (as predicted by SignalPv6.0). Secreted PLA2 enzymes can release free fatty acids from phospholipids (Dennis et al. 2011). These enzymes, which are expressed by human inflammatory cells such as macrophages and T-cells, possess antibacterial and antiviral properties (Triggiani et al. 2006). A second intriguing gene present in this cluster is an *interferon regulatory factor 1 (Irf)-like* gene (*HyS0045.75*). Irf proteins are involved in the immune response in a wide variety of animals (Wang et al. 2024) and were recently identified as being expressed in immune cells in the anthozoans *Nematostella*, *Stylophora*, and *Acropora* (Levy et al. 2021; Cole et al. 2024; Han et al. 2025; Kozlovski et al. 2025). Other differentially expressed genes of this cluster also included an *Alr1-like* gene (*HyS0029.183*) and *Alr2* (*HyS0001.708*) but not *Alr1* (*HyS0031.168*). *Alr1/2* genes encode transmembrane proteins that are vital for self/non-self recognition between *Hydractinia* colonies (Cadavid 2004; Nicotra et al. 2009; Rosa et al. 2010), while the related *Alr1-like and Alr2-like* genes, which are also present in the allorecognition complex of *Hydractinia*, have been hypothesized to play a role in the anti-pathogenic immune response, separate from their role in self/non-self recognition (Nicotra 2022).

We conducted HCR-FISH on feeding polyps to determine the location and morphology of cells in cluster C9 using three specific markers; *HyS0016.300*, an unannotated gene (Fig 6c), and the previously discussed genes *Irf-like* (*HyS0045.75*), and *conodipine-like* (*HyS0053.57*) (Supplementary Fig. S6a-b). Each of these markers were predominantly expressed in cells of the epithelial ectoderm, distributed widely over the polyp body, including within some cells in the tentacles and hypostome (Fig. 6d-f, Supplementary Figs. S6a’-a’’’ and b’-b’’’). In approximately half of the polyps, each marker also showed expression in one or more clusters of cells (upper box in Fig. 6d, higher magnification image in Fig. 6e-e’). The biological significance of these cell clusters is unclear.

The morphology of cells expressing *Hy0016.300* was varied. We obtained high-magnification images of individual cells from whole-mount HCR-FISH samples (Fig. 6g-i’) and performed HCR-FISH on dissociated cells (Fig. 6 j-l’). Some cells were elongated with a central nucleus (Fig. 6g-g’, j-j’, others were round (Fig. 6h-h’, k-k’), while others were irregularly shaped (Fig. 6i-i’, l-l’). Cells that expressed either *Irf-like* or *conodipine-like* showed less morphological diversity and were generally round (Supplementary Fig. S6a’-a’’’ and b’-b’’’). It is unclear if this morphological diversity indicates cells within cluster C9 consist of more than one cell type or whether cells within this cluster change their shape due to biological reasons (e.g., cell migration or phagocytosis).

A recent single-cell analysis in *Hydractinia* also identified a discrete cluster of cells with transcriptomic characteristics similar to cluster C9 in our atlas. These were labeled as conodipine+ or, alternatively, as venomous epithelial cells (Salamanca-Díaz et al. 2025). Using immunofluorescence targeting Alr1 (LOC130635932), the authors report that these cells are widespread in the ectodermal epithelial layer of feeding polyps at the aboral end of the polyp and in the stolon. However, *Alr1* (LOC130635932) is not specifically expressed in the “venomous epithelial cell” cluster in the Díaz *et al*. atlas; instead, it is rather widespread. Furthermore, the feature plot of an *Alr1-like* gene (LOC130635943) in Fig. 5B of that study would not be recognized by the Alr1 antibody used to generate the images in their Fig. 5C. Similarly, *Alr1* (*HyS0031.168*) is also not specifically expressed in cluster C9 in our atlas, but instead is broadly expressed across the entire UMAP. Therefore, the immunofluorescence pattern of Alr1 shown in Salamanca-Díaz et al. (2025) and the HCR-FISH patterns for the genes we used to highlight our C9 cluster (*HyS0016.300, conodipine-like, and Irf-like*) cannot be directly compared or expected to give similar spatial patterns.

Taken together, bioinformatic and marker gene expression analyses have led us to hypothesize that the cells comprising C9 represent a distinct type of ectodermal cell specifically involved in host defense and immunity in *Hydractinia*. These cells might be involved in the identification of pathogens and/or the downstream responses that potentially involves phagocytosis and intracellular digestion. Epithelial cells in *Hydra* (Bosch and David 1986) and anthozoan amoebocytes have been shown to be phagocytic and, in some cases, migratory (Olano and Bigger 2000; Mydlarz et al. 2008; Parisi et al. 2020; Snyder et al. 2021). Ultimately, determining the precise function of the cells that constitute cluster 9 will require further experimentation that is beyond the scope of this study. For example, future experiments could investigate the response of these cells when *Hydractinia* is exposed to pathogenic organisms or other non-self challenges. Additionally, phagocytosis assays could be performed.

### I-cells and Progenitors

I-cells are adult stem cells in *Hydractinia*. They are found throughout the colony, including in feeding polyps, sexual polyps, and the stolon. In feeding polyps, they are located primarily in the epidermal layer and are most dense in a band-like region in the aboral half of the polyp body. They are characterized by their size (7-10 µm), large nuclear-to-cytoplasmic ratio, and high ribosomal content (Plickert et al. 2012). The C3 cluster was annotated as i-cells and progenitors due to the expression of known i-cell markers such as *Piwi1* (*HyS0050.7*), *Myc (HyS0005.84*), and *Nanos1* (*HyS0036.26*) (Plickert et al. 2012). C3 was connected to one of the neural clusters (C7) and three cnidoblast clusters (C6, C15, C16) that are known to differentiate from i-cell precursors (Fig. 1b, Supplementary Fig. S1) (Varley et al. 2023).

To further investigate the i-cell population, we performed a subclustering analysis to determine whether the C3 cluster might contain transcriptionally distinct subpopulations. After subsetting C3 from the larger dataset (Fig. 7a), we re-normalized the data and generated t-SNE plots for visualization (Fig. 7b). Using a similar approach to the one we applied to the full UMAP, we annotated these subclusters using the top differentially expressed genes of each cluster (Supplementary Table S5), combined with literature searches to identify genes that had previously identified functions in *Hydractinia,* other cnidarians, and other animals.

**Figure 7.**
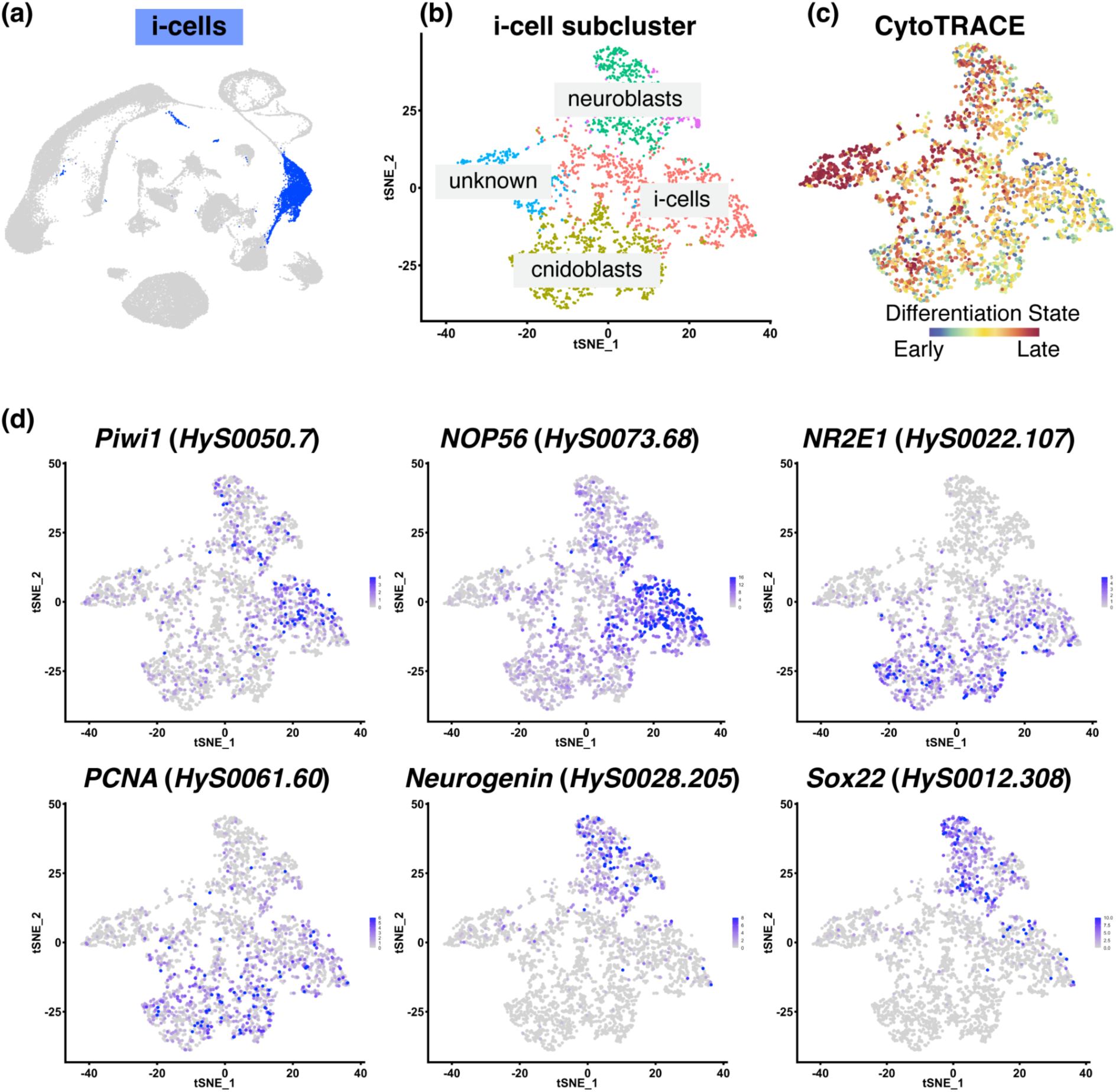
I-cell subcluster analysis. (a) Two-dimensional UMAP of the *Hydractinia* single-cell atlas with cluster 3 (i-cells/progenitor cells) highlighted in blue. (b) The i-cell cluster was selected and reclustered to generate the i-cell subcluster atlas. This subclustering analysis allowed us to designate putative cell states. (c) CytoTRACE analysis was performed on the i-cell subcluster object, revealing the differentiation state from early (blue/green) to late (orange/red). (d) UMAP expression of specific genes that were used to annotate the clusters in (b).

The canonical i-cell marker, *Piwi1*, was predominantly expressed in the largest subcluster (Fig. 7d). Genes that showed differential expression in this cluster also included those involved in ribosome biogenesis, such as *NOP56* (*HyS0073.68*)*, NOP58* (*HyS0155.10*) (Waletich et al. 2024), and *GNL3* (*HyS0059.86*) (Quiroga-Artigas et al. 2022), as well as genes encoding ribosome subunits *RPL38* (*HyS0006.38*), and *RPL23* (*HyS0023.266*) (Fig. 7d, Table S5). Given that *Hydractinia* i-cells have been described as “rich in ribosomes” (Plickert et al. 2012) – a feature shared with mammalian embryonic stem cells that are also known to display elevated ribosomal gene expression (reviewed in Gupta and Santoro 2020) – as well as the high levels of *Piwi1* observed in this cluster, we labeled this subcluster as the true i-cells.

The second largest subcluster expressed known marker genes for cnidoblasts, *NR2E1* (*HyS0022.107*) (Siebert et al. 2019), *Txd12* (*HyS0042.111*) and *Fkbp14* (*HyS0020.22*) (this publication), as well as genes involved in the DNA replication machinery, including *PCNA* (*HyS0061.60*) (Waletich et al. 2024) and *MCM7* (*HyS0009.219*). Cnidocytes are single-use cells that are required to be constantly replenished via cell division from i-cell precursors. Therefore, the presence of specific cnidoblast markers combined with genes involved in cell proliferation led us to label this cluster as cnidoblasts. Next, we annotated two subclusters as neuroblasts, based on expression of neurogenesis genes such as *Neurogenin* (*HyS0028.205*) and *Sox22* (*HyS0012.308*) (Fig. 7d). A final cluster remained unannotated (“unknown”, Fig. 7b) due to having a very short list of differentially expressed genes (Table S5). CytoTRACE analysis of the entire i-cell subcluster showed that the cells of this cluster were more differentiated compared to cells in other subclusters, indicating that it might contain progenitor cells of a yet unidentified cell type or represent a transitory state between two cell types (Table S5, Fig. 7c).

We did not identify a subpopulation of cells in C3 that could serve as progenitors to epithelial or gland cells. These progenitor cells may be present in the unannotated subcluster or exist in very low numbers, making them undetectable at the current clustering resolution. In *Hydra*, it has been shown that, unlike cnidocytes, epithelial and gland cells have a slower turnover rate (David and Campbell 1972; David and Gierer 1974; Bode et al. 1987). The absence of lineage connections between i-cells and these cell types in the atlas presented here, as well as in previous single-cell atlases, (Schnitzler et al. 2024; Salamanca-Díaz et al. 2025), suggests that the same is true for *Hydractinia*.

Genes such as *Piwi1, Myc,* and *Nanos1* have long been used as markers for i-cells and pluripotency in *Hydractinia* (Plickert et al. 2012; Bradshaw et al. 2015). A recent article detailing the migration of a single *Piwi1-GFP^+^* cell and subsequent proliferation and differentiation into all cell types provided direct evidence for their pluripotency (Varley et al. 2023). In the i-cell subclustering analysis detailed here, *Piwi1* expression was predominantly localized to one subcluster; however, it was not exclusive to that subcluster – for example, neuroblast subcluster cells also express *Piwi1* (Fig. 7d). This parallels findings in other highly regenerative organisms, such as planarians, where lineage-primed progenitors (including neural-specified neoblasts) retained pluripotency (Fincher et al. 2018; Raz et al. 2021). These observations suggest that, while *Piwi1* is a useful marker, it alone cannot fully define adult stem cell identity. Instead, our results support a model where adult stem cells (ASCs) likely comprise a heterogeneous population that includes both undifferentiated and lineage-primed progenitors, with dynamic gene expression that adapts to cellular contexts (Rinkevich et al. 2022).

### Conclusions

Here, we present an updated and spatially validated somatic single-cell atlas for *Hydractinia* feeding polyps and stolons. By integrating a previously published live-cell dataset with newly generated fixed-cell datasets, we comprehensively recapitulate all known cell lineages—except the germline—while expanding the current understanding of *Hydractinia*’s cellular landscape. Key discoveries include: A novel neural subtype (Neurons B) primarily found in tentacles, two spatially distinct gland cell populations, and a putative immune cell type. We provide the first complete cnidogenesis trajectory for *Hydractinia*, validating the expression of key markers along this pathway that splits into two distinct terminal endpoints representing desmonemes and euryteles. This trajectory enables comparative analysis with existing trajectories from *Hydra*, *Nematostella*, and *Clytia*, offering new insights into the evolution of this specialized cnidarian cell type. We also reclustered the somatic i-cell population and successfully identified distinct subclusters, assigning putative cell states to each. This analysis uncovered a population of *bona fide* i-cells, along with early progenitor populations for specific cell types that will be useful in exploring the transcriptional dynamics governing stem cells and early progenitors. Notably, the absence of shared neuroglandular progenitors in our atlas aligns with findings in *Clytia* but contrasts with *Hydra* and *Nematostella*; further studies incorporating young or regenerating polyps could clarify the relationship between neurons and gland cells in *Hydractinia*. Future work could also focus on characterizing the putative immune cell type through pathogen challenges and phagocytosis assays. This updated atlas provides a wealth of new data, promising candidate genes for creating transgenic reporter lines, and a foundation for deeper characterization of specific cell types and cell states through targeted functional investigations.

## Materials and Methods

### Animal Husbandry

Adult *Hydractinia symbiolongicarpus* colonies (291-10, male) were maintained at the University of Florida, Whitney Laboratory for Marine Bioscience. Colonies were grown on glass microscope slides and cultured in 38 L tanks filled with artificial seawater (Coral Pro Salt, Red Sea) at 30 ppt and kept at 18–20 °C under a 10 h/14 h light/dark regime. Animals were fed five times a week with 3-day-old brine SEP-Art *Artemia* nauplii (INVE Aquaculture), which were enriched two times with S.presso (SELCO) the day before colony feeding.

### Single-cell Dissociation

Protocol was adapted from a previous *Hydractinia* single-cell study (Schnitzler et al. 2024). Twenty feeding polyps (gastrozooids) and their surrounding stolonal tissue were removed from the colonies and washed three times in calcium- and magnesium-free seawater (CMFSW: 450 mM NaCl, 9 mM KCl, 30 mM Na_2_SO_4_, 2.5 mM NaHCO_3_, 10 mM Tris-HCl, 2.5 mM EGTA, 25 mM HEPES). The polyps were then placed in 300 μL 1% pronase (Santa Cruz Biotechnology, catalog # sc-264144) in CMFSW for 1.5 h on a rocker at room temperature. Every 15 min the tube was gently mixed by inverting. Once the tissue was fully dissociated, the cell suspension was filtered using 70 μM Flowmi tip filter (Bel-Art, catalog # H13680-0070) into a 2 ml DNA LoBind tube (Eppendorf, catalog # 022431048). The sample was centrifuged at 300 g for 5 min at 4 °C. Supernatant was gently removed while leaving about 50 μL at the bottom of the tube, then 500 μL CMFSW was added to resuspend the cell pellet. The sample was centrifuged again at the same settings above, supernatant removed and resuspended in 200 μL CMFSW. Cells concentrations were determined using a hemocytometer.

### Cell Fixation

Dissociated cells were then immediately fixed using two different fixatives: 800 μL ice-chilled 100% methanol (Sigma-Aldrich) or ACME solution (13:3:2:2 ratio of DNase/RNase-free distilled water, methanol, glacial acetic acid, and glycerol) (García-Castro et al. 2021). Two samples were prepared using the methanol method and eight samples were prepared using the ACME method. Both fixatives were added dropwise to the cell sample. Fixed cells were then transferred to a -20 °C freezer for storage.

### Single-cell RNA sequencing

Fixed single cell samples were diluted to 1,000 cells/μL and shipped on dry ice to the National Institute of Health Intramural Sequencing Center (Bethesda, MD). Cells were thawed, spun and resuspended but cell counts were not obtained again, and the samples were loaded into the 10X Genomics platform for encapsulation with the capture target of 6,000-9,000 cells per sample (Table S1). Sequencing libraries were prepared according to the standard 10X Genomics V3 chemistry protocol. The cDNA libraries were pooled and sequenced as 150bp paired end reads and single indexed on an Illumina NovaSeq6000 with 63 million projected clusters per sample. Raw sequencing data were processed with the CellRanger v7 pipeline (10X Genomics), using default parameters and expected recovery of 6,000-9,000 cells for each respective library.

### Data Processing and Bioinformatic Analyses

After preliminary QC in CellRanger, one of the ACME-fixed samples was discarded due to poor quality. In addition to the two methanol-fixed samples and seven remaining ACME-fixed samples, we also included a previously published single-cell dataset of *Hydractinia symbiolongicarpus* in our analyses (Schnitzler et al. 2024). To focus on somatic cell lineages, cells expressing sperm-related markers (*HyS0027.170, HyS0070.46, HyS4524.1, HyS0007.253, HyS0001.110*) were subsequently removed from the previous dataset. All count matrices were individually processed and cleaned using Seurat v5.2.1 (Stuart et al. 2019; Hao et al. 2021) in R. In short, potential cell multiplets were removed by using a library-specific cutoff for aberrantly high UMI counts and gene counts (for detailed sample processing, see code in “Data Availability”). After the initial filtering, we ran the dataset through a standard Seurat analysis pipeline using the default parameters unless otherwise specified as follows: data were normalized and variable features were selected by running “*SCTransform*”, vst.flavor = “v2” (Choudhary and Satija 2022). The top 50 principal components were calculated with the *RunPCA* function. Clustering was performed by running the *FindNeighbors* function with dims= 1:15. This was followed by running *FindClusters* with resolution=0.5. Nonlinear dimensionality reduction was performed to represent the data in a 2D space using Uniform Manifold Approximation and Projection (UMAP) (McInnes et al. 2018).

Given that all our samples are predicted to include most cell types of the animal, and no significant technical variation was expected, we chose canonical correlation analysis (CCA) to integrate different datasets (Butler et al. 2018). We selected 3,000 genes by running *SelectIntegrationFeatures* and integrated datasets by running *IntegrateData*, normalization.method = “SCT”. The integrated dataset was then processed using the standard Seurat pipeline above, with 50 principal components, dims=1:20 in clustering and resolution=0.3 in UMAP. Differential expression (DE) analyses were identified with the *FindAllMarkers* function, with min.pct=0.1, min.diff.pct=0.5, logfc.threshold=1, using the “RNA” assay. Clusters were annotated based on the DE gene list and known cell type markers (Schnitzler et al. 2024). A list of genes used to annotate all the clusters and their expression in the single-cell atlas can be found in Supplementary Table S4.

### Cnidogenesis Single-Cell Atlas

The updated *Hydractinia* single-cell atlas was subset to create a cnidogenesis atlas using R (v4.4.0) and the Seurat v5.1.0 package (Hao et al. 2021). I-cells and cnidocyte clusters (C1, C2, C3, C4, C6, C8, C15, and C16) were extracted from the whole Seurat single-cell object to create a cnidogenesis specific object. Integration anchors were calculated to reduce batch effects from the different single-cell libraries. Finally, dimensionality reduction and clustering analysis were performed to generate the final version of the *Hydractinia* cnidogenesis single-cell atlas. In order to align the orientation of the cnidogenesis differentiation trajectory in the whole atlas with the cnidogenesis atlas trajectory, the x-axis of the cnidogenesis atlas dimensionality reduction plots (e.g., FeaturePlot, DimPlot) were reversed using scale_x_reverse() in the ggplot2 package (Wickham, H. 2016).

### CytoTRACE Differentiation State Analysis

The R package cytoTRACE v0.3.3 was used to predict the differentiation state of cells in the *Hydractinia* cnidogenesis single-cell atlas and the i-cell subcluster atlas (Gulati et al. 2020). Differentiation scores for each cell were computed, added to the metadata of the relevant Seurat object, and then visualized using the Seurat FeaturePlot function. The cytoTRACE differentiation scores were inverted so that less differentiated cells had lower scores, and more differentiated cells had higher scores.

### Monocle3 Trajectory and Pseudotime Analysis

The cnidogenesis Seurat object was converted into a Monocle3 (v1.3.7) cell data object (Cao et al. 2019) and original Seurat PCA and UMAP embeddings were manually added to the metadata. The cellular trajectory was predicted using the learn_graph function and pseudotime was estimated by manually selecting the i-cell cluster as the root.

### Gene Ontology

Gene ontology enrichment analysis was performed and visualized using the R package topGO v.2.54.0 (Alexa and Rahnenfuhrer). The corresponding GO term accessions were retrieved from a customized text file (supplement “Hsym_v1.0_GO_terms.out”) for *Hydractinia*. Genes of interest were the differentially expressed genes from each cluster (Table S4). Enrichment tests were performed using the arguments algorithm=‘classic’, statistic=‘fisher’.

### Neuropeptide Predictions

Putative neuropeptides were predicted based on the method in (Chari et al. 2021). *Hydractinia* predicted proteins were downloaded from the genome project portal (https://research.nhgri.nih.gov/hydractinia/) and screened for the presence of a signal peptide using SignalP v6.0 (Teufel et al. 2022). Proteins deemed to be transmembrane proteins were removed, based on predictions from SignalP v4.0. A custom Perl script was then implemented to screen the remaining proteins that possessed a signal peptide for the presence of one or more neuropeptide cleavage sites (G[KR][KRED]). When more than one site was present, the 6 residues immediately N-terminal to this cleavage site were compared with each other. Putative neuropeptides were ranked according to a normalized score, where the sum of identical amino acids at each position for each 6 AA motif were divided by the number of motifs present in a protein. Expression profiles of all putative neuropeptides in the single cell atlas were then investigated. Those that had a normalized score of 1 or more that were also expressed predominantly in one or both neural clusters (clusters 7 and/or 14) were selected. This shortlist was further refined by manually comparing the 6 amino acid motifs within each protein to each other to ensure similarity. Finally, a list of 12 putative neuron-specific neuropeptides was generated. UMAP embeddings of each of these genes are shown as Supplementary Fig. S4, and amino acid sequences shown as Table S2.

### I-cell/progenitors Subcluster Analysis

Cluster 3 was subjected to further subclustering analysis to investigate potential cell subpopulations. The three datasets were SCTransformed (vst.flavor = “v2”) individually as mentioned above and reintegrated using CCA with 3,000 features (genes). The integrated dataset was then processed using the standard Seurat pipeline, with 25 principal components, dims=1:25 in clustering and resolution=0.2 in t-SNE projection. Differential expression (DE) analyses were identified with the *FindAllMarkers* function, with min.pct=0.3, logfc.threshold=1, using the “RNA” assay.

### HCR fluorescent *in situ* hybridization (HCR-FISH)

For each cell cluster, the top differentially expressed marker genes were examined to determine their suitability for HCR. Genes that were particularly specific to the cluster of interest, had a very high level of expression as determined by the number of transcripts present, and where eight or more probe pairs could be designed were chosen for spatial analysis using HCR-FISH. The number of probe pairs was limited to 40 when necessary. DNA probe sets were designed using the Ӧzpolat Lab probe generator (https://github.com/rwnull/insitu_probe_generator) (Kuehn et al. 2022). The sequences generated by the algorithm were used to order DNA oPools™ Oligos from Integrated DNA Technologies (IDT), which were resuspended in nuclease-free H_2_O to a final concentration of 1 pmol/μL. All buffers and hairpin amplifiers were ordered from Molecular Instruments, Inc. The HCR-FISH protocol for *Hydractinia* was based on published methodology (Choi et al. 2018). Adult feeding polyps dissected from the stolon mat and whole juvenile colonies were relaxed in 4% MgCl_2_ 1:1 filtered seawater (FSW):H2O before being fixed in 4% paraformaldehyde (PFA) in 1x PBS + 0.1%

Tween-20 (PTw) for 1-2 hours at 4oC. Samples were then dehydrated in increasing concentrations of methanol in PTw (25%, 50%, 75%, 100%) and stored at −20°C for at least 2 hours. Following rehydration in a reverse methanol:PTw series (100%, 75%, 50%, 25%), samples were washed several times in PTw, before incubation in a solution of 50% PTw:50% probe hybridization buffer for 15 minutes at room temperature. Prehybridization was conducted for 1 hour at 37°C in 100% probe hybridization buffer. Following the -one-hour prehybridization step, gene-specific probe sets were added to a final concentration of 20-40 nM, depending on the gene, and were generally hybridized for 16-24 hours at 37°C. For two genes (HyS0045.75 and HyS0053.57), we found that the signal was improved by hybridization of probes for 6 days. After hybridization, prewarmed wash buffer was used to wash samples 4 x 15 minutes at 37°C, followed by 3 x 5-minute washes with 5x SSCT (5x SSC, 0.1% Tween-20) at room temperature. Samples were then incubated in an amplification buffer for 30 minutes at room temperature. During this step, hairpins were prepared by adding 6 pmol of each hairpin (h1 and h2) into separate 0.5mL tubes (the hairpin/fluorophore combination depended on the probe sets used) and heated to 95°C for 90 seconds. Hairpins were then cooled to room temperature in the dark for 30 minutes. Finally, hairpin pairs were combined, and the appropriate volume of amplification buffer was added to create a ‘hairpin solution’ with a final volume of 100 μL. The pre-amplification solution was removed from samples and the appropriate ‘hairpin solution’ added to each tube. Samples were incubated overnight at room temperature in the dark. Samples were washed in 5x SSCT for 2 x 5 mins, 2 x 30 mins and finally 1 x 5 mins. Hoechst 33342 (ThermoFisher H1399) was included in one of the 30-minute wash steps at a final concentration of 10 μg/mL to stain nuclei. Finally, samples were mounted in 70% ultrapure glycerol:PBS before confocal imaging. Negative controls were included for all hairpins used, where the procedure was followed as normal, however probe sets were not added to the hybridization solution. Images of negative controls were captured using the same confocal settings used for experimental samples to ensure background fluorescence was not mimicking real signal. The complete list of probe sets and associated initiators can be found in Table S6.

### HCR-FISH on Dissociated Cells

Approximately 40 adult feeding polyps were relaxed in 4% MgCl_2_ 1:1 filtered seawater (FSW):H_2_O for at least 15 minutes before being dissected from the stolon mat. Animals were decapitated and ‘heads’ and ‘bodies’ dissociated separately. Polyps were then washed two times in ACME solution in FSW (13:3:2:2 = FSW:methanol:acetic acid:glycerol), before being washed two times in ACME solution in diH_2_O (13:3:2:2 = diH_2_O:methanol:acetic acid:glycerol). Polyps were dissociated by vigorously pipetting solution up and down in 1 mL of ACME solution in diH_2_O for several minutes. An ImmEdge^®^ Hydrophobic Barrier PAP Pen was used to draw a circle on a SuperFrost slide (Cat. 12-550-15); 200 μL of dissociated cells were pipetted into the center of the circle and cells were left to settle overnight. HCR-FISH and Hoechst nuclei staining was performed on slides as above before imaging.

### Fluorescent *in situ* hybridization (FISH)

Adult *Hydractinia* colonies were placed in a solution of 4% MgCl_2_ in distilled water:filtered seawater (FSW) (1:1) for 10-15 minutes, before feeding polyps were cut from the stolon mat. Polyps were fixed for 90 seconds in an ice-cold solution of 0.2% glutaraldehyde, 4% paraformaldehyde (PFA) and 0.1%Tween-20 in FSW, followed by fixation in an ice-cold solution of 4% PFA and 0.1% Tween-20 in FSW for 90 minutes at 4°C. Following fixation, samples were washed multiple times with ice-cold DEPC-PTw (1x phosphate-buffered saline (PBS) with 0.1% Tween20 in DEPC-treated H_2_O) before being dehydrated with increasing concentrations of methanol in DEPC-PTw (25%, 50%, 75% and 100%). Digoxigenin (DIG)-labeled riboprobes were generated with the SP6 or T7 MEGAscript kit (catalog #AM1334, #AM1330, Ambion, Inc., Austin, TX, USA). Immediately prior to *in situ* hybridization, samples were rehydrated with decreasing concentrations of methanol in DEPC-PTw, followed by several washes in DEPC-PTw. Samples were then washed for five minutes each in 1% triethylamine in DEPC-PTw (TEA), 0.6% acetic anhydride in TEA, and 1.2% acetic anhydride in TEA, followed by several washes in DEPC-PTw. Samples were pre-hybridized for 4 hours at 55°C in hybridization buffer (4M urea, 0.1 mg/ml yeast tRNA, 0.05 mg/ml Heparin, 5x SCC pH7.0, 0.1% Tween20, 1% SDS in DEPC-treated H_2_O). Riboprobes were diluted to a concentration of 0.5 ng/μL in hybridization buffer and heated to 90°C for 10 minutes before being added to samples and incubated for approximately 40 hours at 55°C. Following hybridization, unbound probe was removed in a series of washes; hybridization buffer at 55°C for 40 minutes and then decreasing hybridization buffer concentrations in 2x SSC at 55°C, followed by washes with decreasing concentrations of 0.2x SSC in PTw at room temperature (RT). Endogenous peroxidase activity was quenched by two 30-minute washes in 3% hydrogen peroxide (H_2_O_2_), followed by further washes in PTw. Two 10-minute washes in maleic acid buffer (MAB, 100mM Maleic acid, 150mM NaCl, pH7.5) were then conducted. Samples were blocked for one hour in blocking buffer (Sigma-Aldrich, Cat. #11096176001 diluted 1:10 in MAB). Bound DIG-labeled riboprobe was detected by incubating samples overnight in 1:1500 dilution of Anti-DIG-POD antibody (Roche, Cat. # 11207733910) at 4°C. Unbound antibody was removed by washing samples several times at room temperature in MABX (MAB containing 0.1% Triton X-100). Samples were then incubated in tyramide development solution (2% Dextran sulfate, 0.0015% hydrogen peroxide, 0.2mg/ml Iodophenol, 1:100 Alexa Fluor 594 Tyramide Reagent (Thermo Scientific, Cat. # B40957) in PTw for eight minutes and then washed several times in PTw. Nuclei were stained using Hoechst dye 33342 (ThermoFisher, Cat. # H1399).

### Microscopy and Image Analysis

All samples were imaged with a Zeiss LSM 710 confocal microscope (Zeiss, Gottingen, Germany), and Z-stack projections were generated using Fiji (Schindelin 2012). All figures were created in Adobe Photoshop (version 25.12.0) or Adobe Illustrator (version 29.1).

## Supplementary Materials

Supplementary_Figures_S1_to_S6.pdf

Supplementary_Tables_S1_to_S6.xlsx

## Acknowledgements

We would like to thank the following individuals for their assistance throughout the course of this study: Alice Young, Jim Thomas, Betty Barnabas, and many others at the NIH Intramural Sequencing Center (NISC) for scRNA-seq library construction, sequencing, advice, and support. We also thank Joe Ryan for bioinformatic and computational support and advice, as well as the members of the Schnitzler lab for many thoughtful discussions and their support.

## Data Availability

Raw sequence data for this study were deposited in NCBI under BioProject ID: PRJNA1263849 with SRA accession numbers: SRR33665854-SRR33665862. All scripts and processed data are available via GitHub: https://github.com/sjwu571/HyS_scRNAseq and Zenodo: https://zenodo.org/uploads/15151309?token=eyJhbGciOiJIUzUxMiJ9.eyJpZCI6ImQ2MjJjODU[…]0bKZQ9RirF3lHeMn8dp80p-ta4AJUo8Q5eDumjz_ijyb48EJCeg_uYASl6ovJBw. A single-cell browser is publicly available at https://sjwu571.shinyapps.io/hys-umap/. Gene specific information for all *Hydractinia* gene IDs discussed in the manuscript can be found at the *Hydractinia* Genome Project Portal: https://research.nhgri.nih.gov/hydractinia/.

