## Supplementary Figures S1 to S6 for "An updated and spatially validated somatic single-cell atlas of *Hydractinia symbiolongicarpus*"

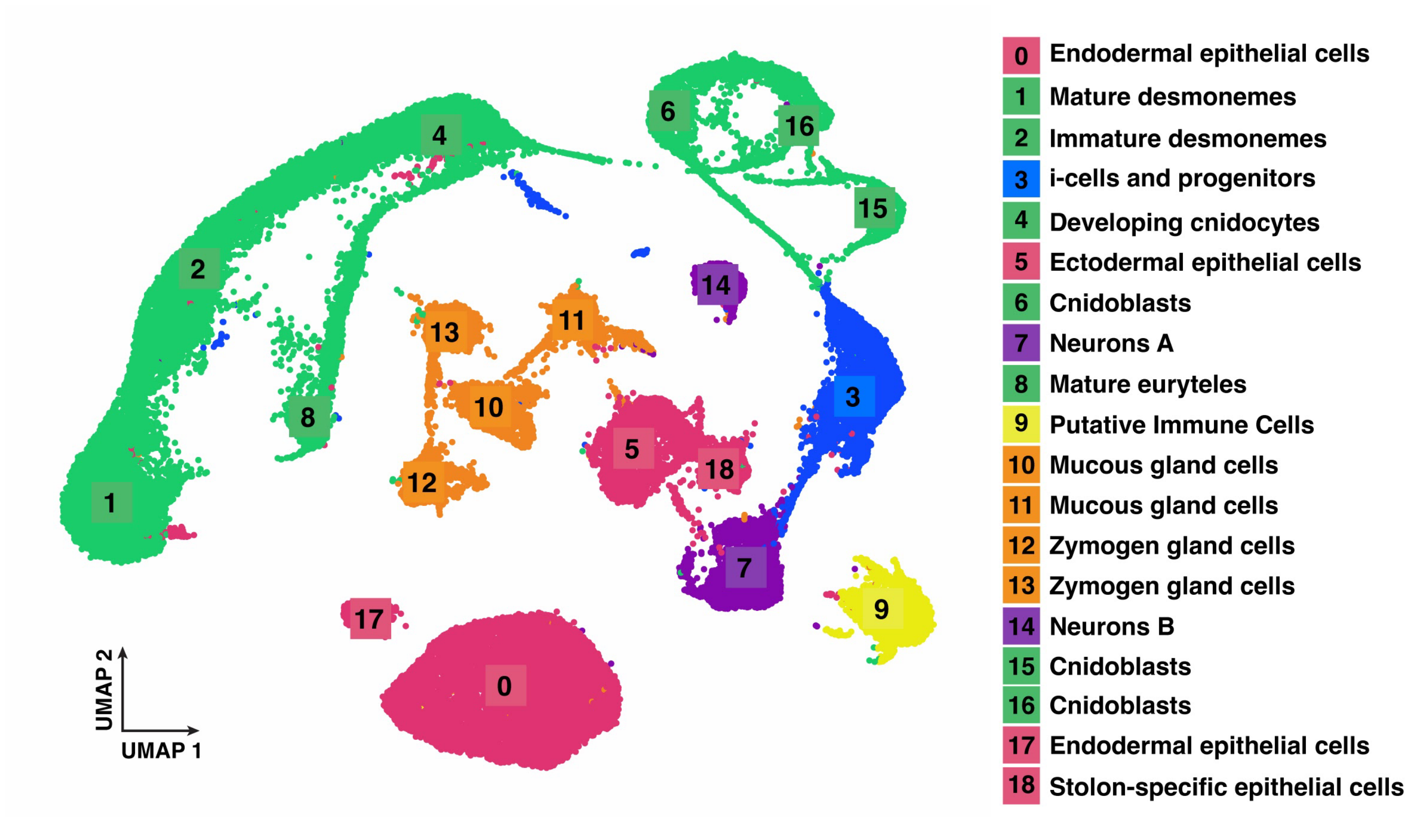


**Supplementary Figure S1. Detailed view of the updated *Hydractinia* single cell atlas.** Two-dimensional UMAP of the updated *Hydractinia* feeding polyp single cell atlas, with all 19 cell states and cell types labelled. Cluster numbers on the UMAP correspond to their hypothesized identity shown in the legend to the right.

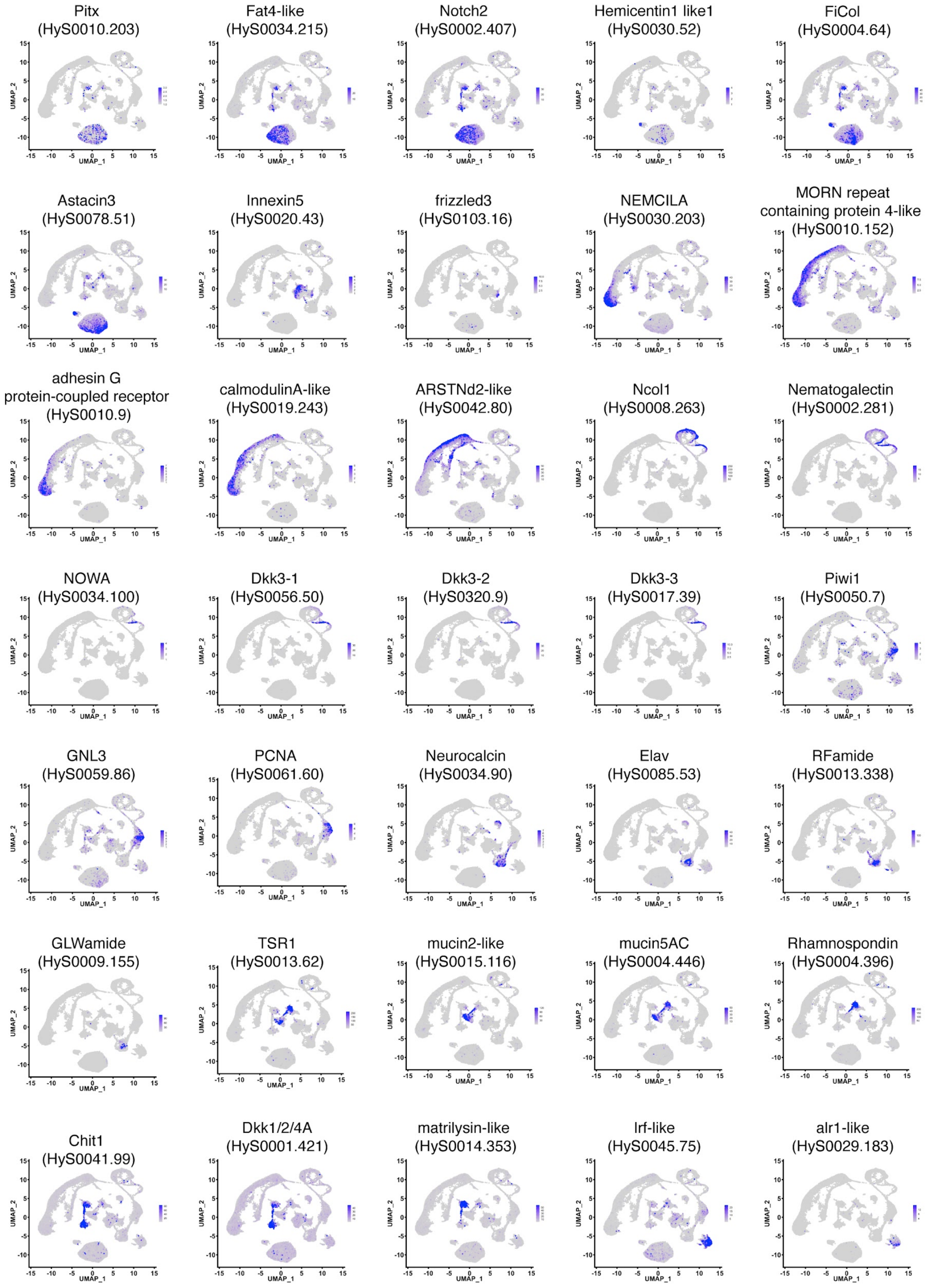


**Supplementary Figure S2. Selected markers used for cluster annotation for the *Hydractinia* single cell atlas.**

The UMAP with labelled clusters is shown in Figure 1b and Supplementary Figure S1.

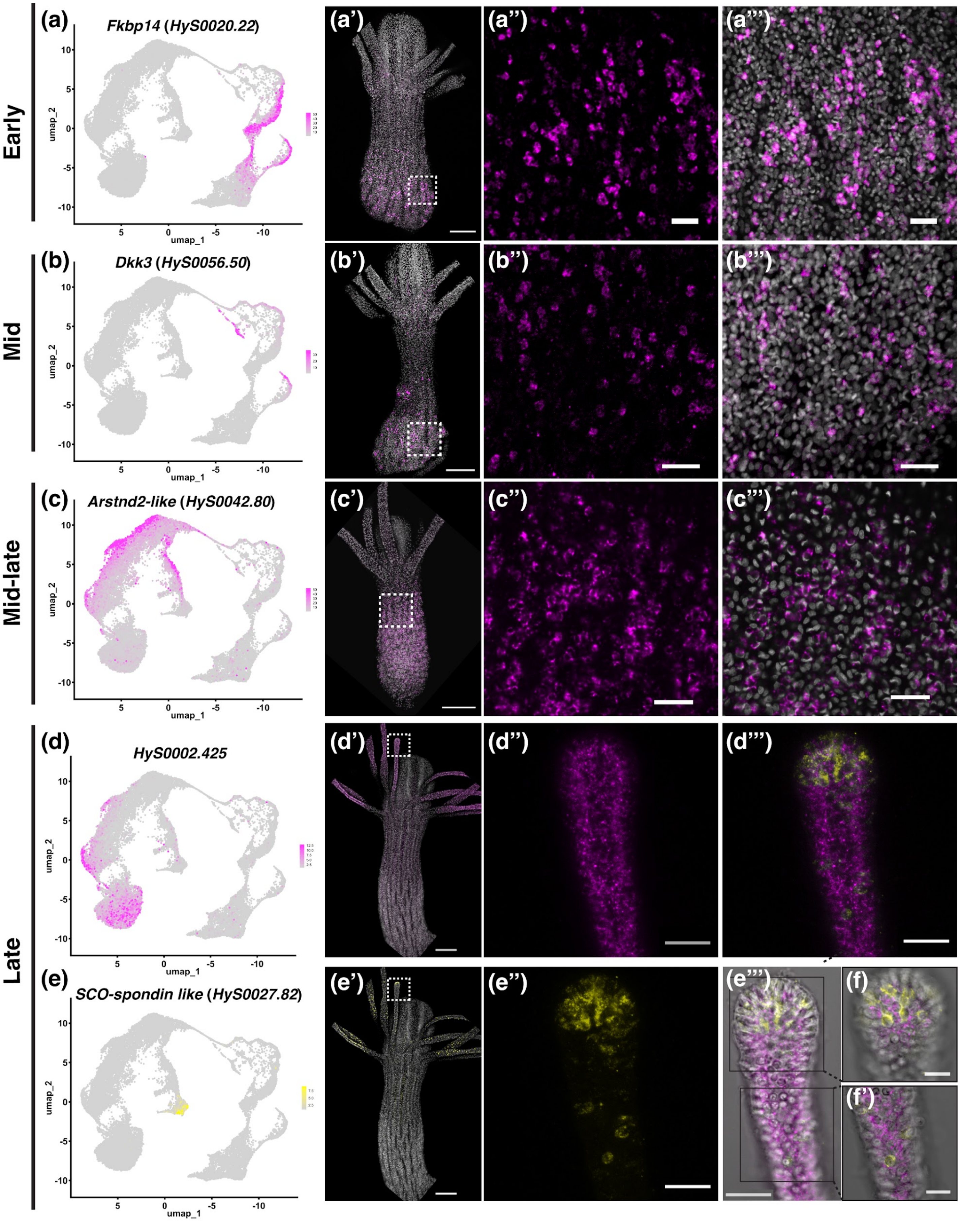
Early
Mid
Mid-late
Late
(a)
Fkbp14 (HyS0020.22)
(a')
(a'')
(a''')
(b)
Dkk3 (HyS0056.50)
(b')
(b'')
(b''')
(c)
Arstnd2-like (HyS0042.80)
(c')
(c'')
(c''')
(d)
HyS0002.425
(d')
(d'')
(d''')
(e)
SCO-spondin like (HyS0027.82)
(e')
(e'')
(e''')
(f)
(f')
umap_1
umap_2

**Supplementary Figure S3. Mapping the transcriptional transition from i-cell to mature cnidocyte.** Left-most column shows the expression of a particular gene in the cnidogenesis atlas, while panels to the right show expression of that gene *in situ* in an adult feeding polyp. Dotted white boxes in (a'), (b'), (c'), (d') and (e') show region of higher magnification shown in (a")-(a'''), (b")-(b'''), (c")-(c'''), (d")-(d''') and (e")-(e''') respectively. Gene expression is shown in magenta or yellow and nuclei are shown in grey. *HyS0020.22* (*Fkbp14*) is a marker of early cnidogenesis (a)-(a'''), *HyS0056.50* (*Dkk3*) is a marker of mid-cnidogenesis (b)-(b'''), *HyS0042.80* (*Arstnd2-like*) is a marker of mid-late cnidogenesis and *HyS0002.425* and *HyS0027.82* are markers of late cnidogenesis (d)-(f'). The same feeding polyp is shown in all panels (d')-(f'). Images in (e''') are a merge of *HyS0002.425* and *HyS0027.82* and brightfield image to show expression surrounding capsules of mature cnidocytes. Black boxes in (e''') shows areas shown for higher magnification images in (f) (upper box) and (f') lower box. Scale bars: 100 μm in (a')-(e'), 10 μm in (f) and (f'), and 20 μm in all other panels.

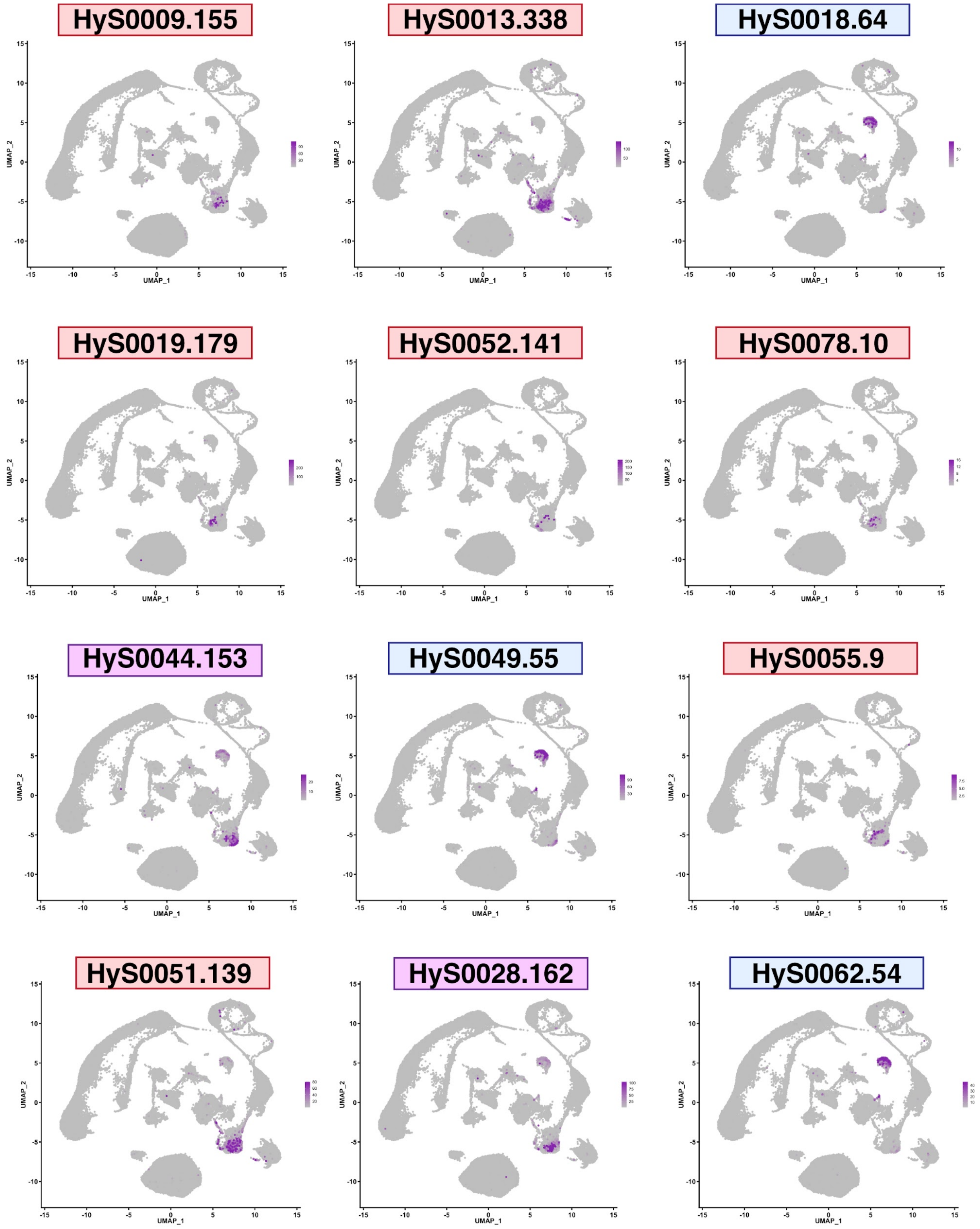


**Supplementary Figure S4. Expression of putative neuropeptides in the neural clusters of the *Hydractinia* single cell atlas.** UMAPs showing expression of predicted neuropeptides that are expressed in cluster 7 (red), cluster 14 (blue) or both clusters (purple).

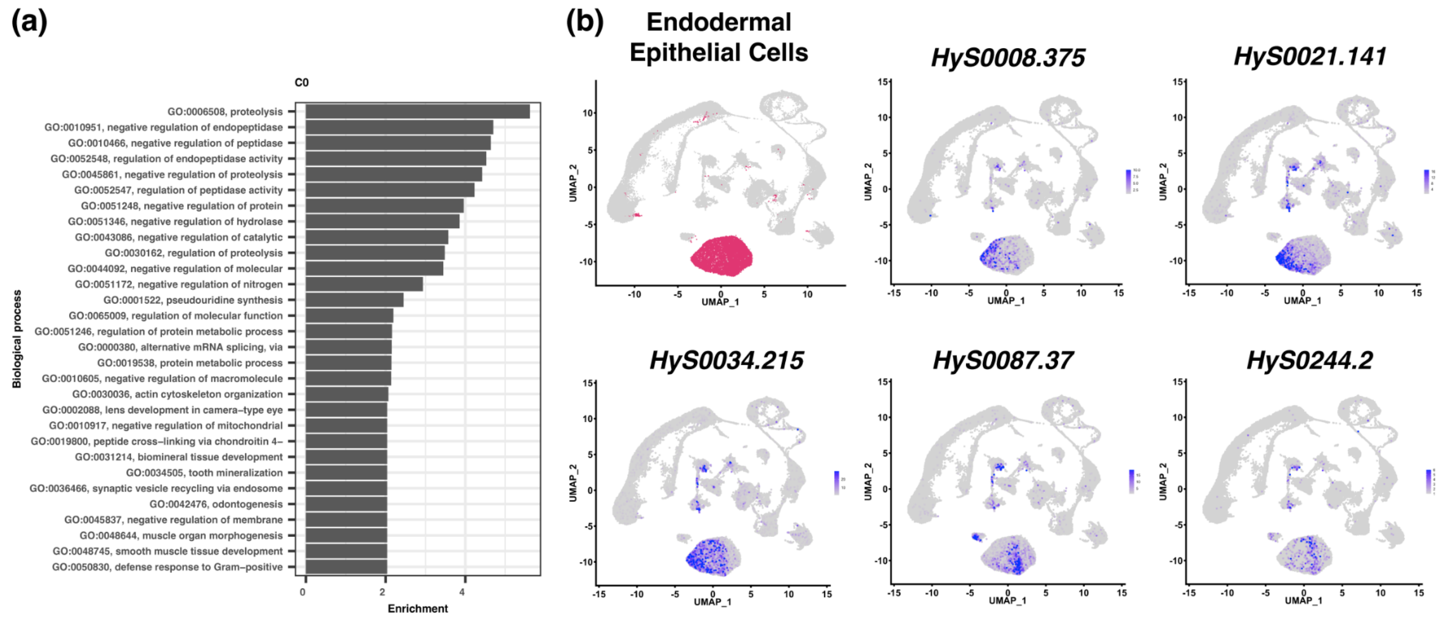


**Supplementary Figure S5. Expression of genes in the endodermal epithelial cell cluster (cluster 0) that show gradient expression across the cluster.** a) Gene ontology enrichment analysis (GO term analysis) of cluster C0. b) UMAPs showing expression of 5 genes that are expressed in cluster 0, with varied expression from left-to-right across the cluster.

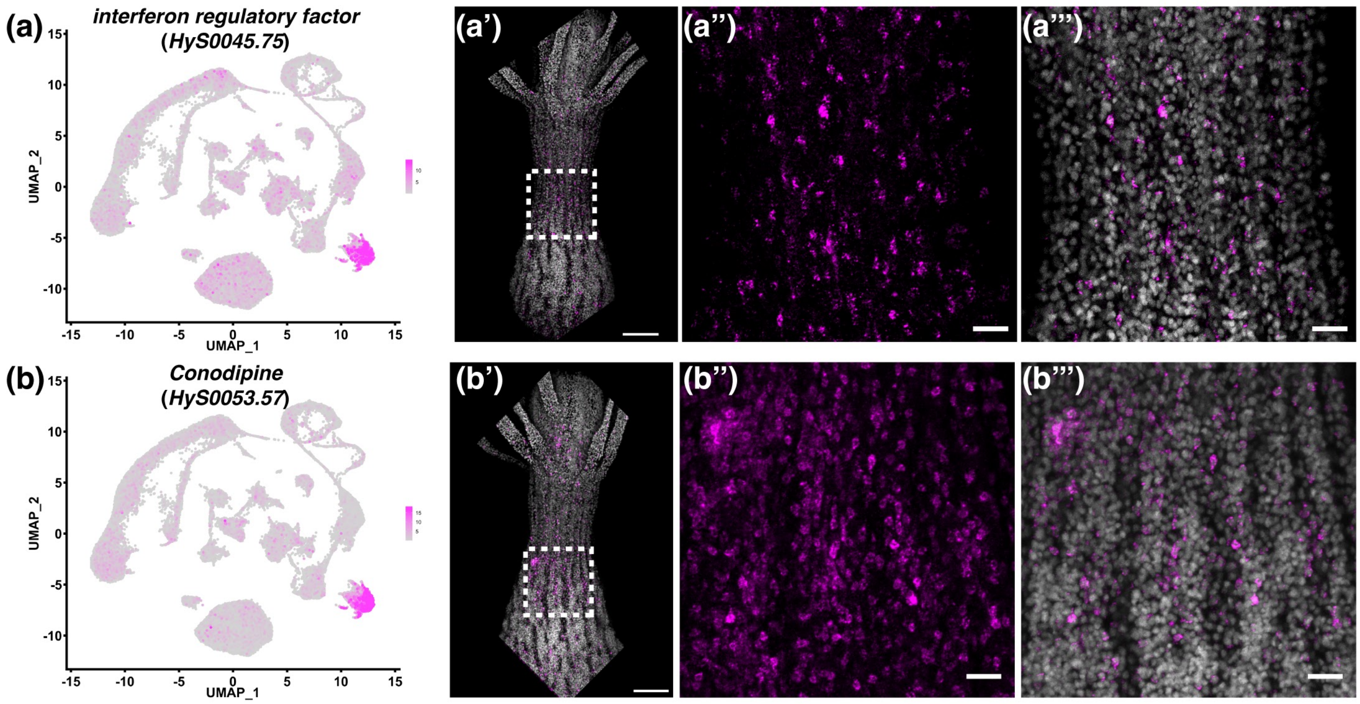


**Supplementary Figure S6. Expression analysis of markers expressed in cell cluster 9, putative immune cells.** (a-a''') UMAP expression of *HyS0045.75* shown in magenta (a) and HCR-FISH of *HyS0045.75* in an adult feeding polyp (a'')-(a'''). (b)-(b''') UMAP expression *HyS0053.57* shown in magenta (b) and HCR-FISH of *HyS0053.57* in an adult feeding polyp (b'')-(b'''). Dotted white box in (a') and (b') shows the area of higher magnification in panels (a'') - (a''') and (b'')-(b'''), respectively. Gene expression is shown in magenta and nuclei are shown in grey. Scale bars: 100 µm in (a') and (b'), 20 µm in all other panels.
